# Real-time single-cell characterization of the eukaryotic transcription cycle reveals correlations between RNA initiation, elongation, and cleavage

**DOI:** 10.1101/2020.08.29.273474

**Authors:** Jonathan Liu, Donald Hansen, Elizabeth Eck, Yang Joon Kim, Meghan Turner, Simon Alamos, Hernan G. Garcia

## Abstract

The eukaryotic transcription cycle consists of three main steps: initiation, elongation, and cleavage of the nascent RNA transcript. Although each of these steps can be regulated as well as coupled with each other, their *in vivo* dissection has remained challenging because available experimental readouts lack sufficient spatiotemporal resolution to separate the contributions from each of these steps. Here, we describe a novel application of Bayesian inference techniques to simultaneously infer the effective parameters of the transcription cycle in real time and at the single-cell level using a two-color MS2/PP7 reporter gene and the developing fruit fly embryo as a case study. Our method enables detailed investigations into cell-to-cell variability in transcription-cycle parameters as well as single-cell correlations between these parameters. These measurements, combined with theoretical modeling, suggest a substantial variability in the elongation rate of individual RNA polymerase molecules. We further illustrate the power of this technique by uncovering a novel mechanistic connection between RNA polymerase density and nascent RNA cleavage efficiency. Thus, our approach makes it possible to shed light on the regulatory mechanisms in play during each step of the transcription cycle in individual, living cells at high spatiotemporal resolution.

## Introduction

The eukaryotic transcription cycle consists of three main steps: initiation, elongation, and cleavage of the nascent RNA transcript (Fig. 1A; ***Alberts (2015***)). Crucially, each of these three steps can be controlled to regulate transcriptional activity. For example, binding of transcription factors to enhancers dictates initiation rates (***Spitz and Furlong, 2012)***, modulation of elongation rates helps determine splicing efficiency (***De La Mata et al., 2003)***, and regulation of cleavage controls aspects of 3’ processing such as alternative polyadenylation (***Tian and Manley, 2016***).

**Figure 1.**
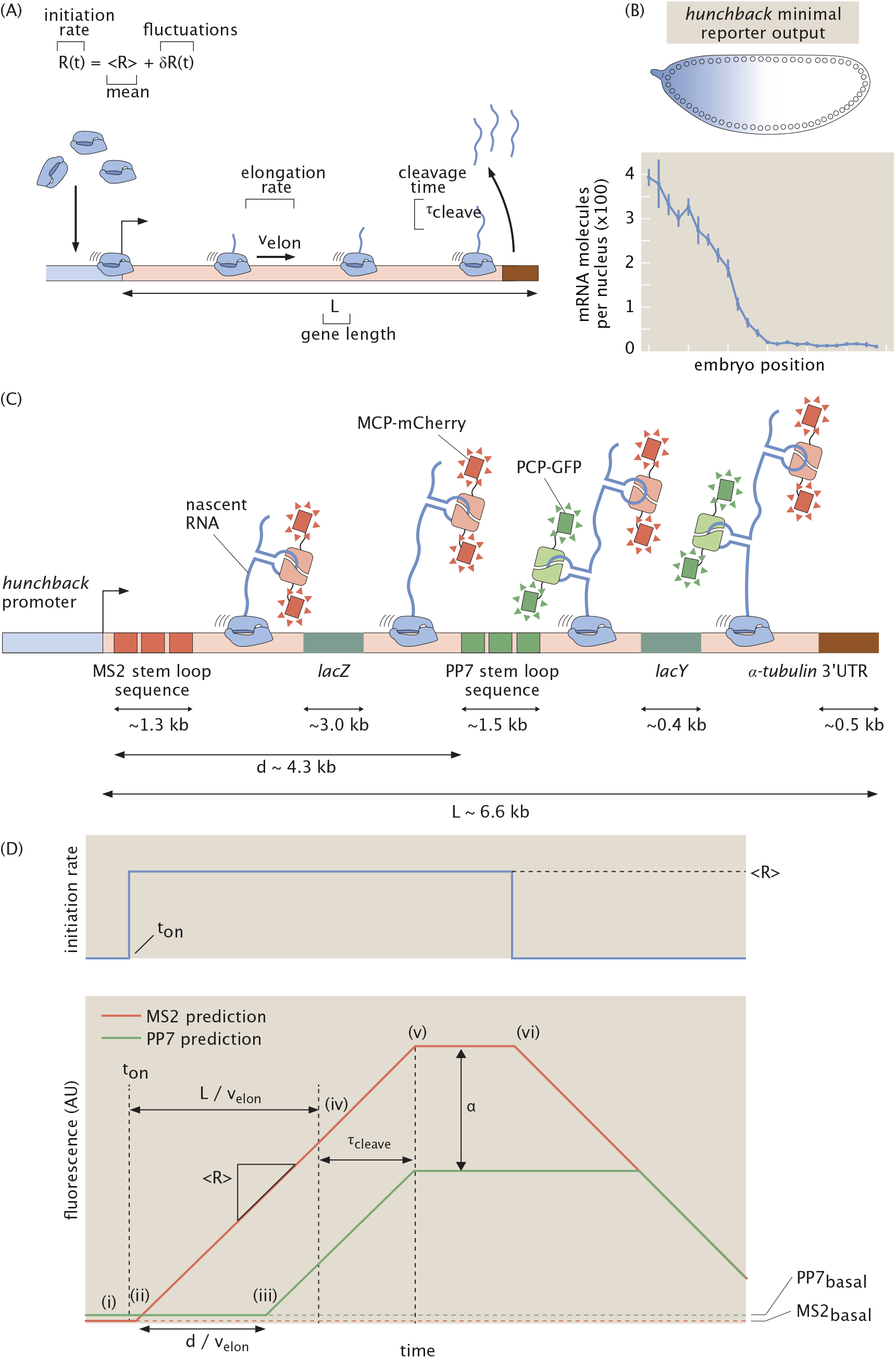
Theoretical model of the transcription cycle and experimental setup. (A) Simple model of the transcription cycle, incorporating nascent RNA initiation, elongation, and cleavage. (B) The reporter construct, which is driven by the *hunchback* P2 minimal enhancer and promoter, is expressed in a step-like fashion along the anterior-posterior axis of the fruit fly embryo. (C) Transcription of the stem loops results in fluorescent puncta with the 5’ mCherry signal appearing before the signal from 3’ GFP. Only one stem loop per fluorophore is shown for clarity, but the actual construct contains 24 repeats of each stem loop. (D, top) Relationship between fluorescence trace profiles and model parameters for an initiation rate consisting of a pulse of constant magnitude ⟨*R*⟩. (D, bottom, i) At first, the zero initiation rate results in no fluorescence other than the basal levels *MS*2_*basal*_ and *PP*7_*basal*_ (red and green dashed lines). (ii) When initiation commences at time *t*_*on*_, RNAP molecules load onto the promoter and elongation of nascent transcripts occurs, resulting in a constant increase in the MS2 signal (red curve). (iii) After time 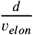, the first RNAP molecules reach the PP7 stem loops and the PP7 signal also increases at a constant rate. (iv) After time 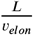, the first RNAP molecules reach the end of the gene, and (v) after the cleavage time *τ*_*cleave*_, these first nascent transcripts are cleaved. The subsequent loss of fluorescence is balanced by the addition of new nascent transcripts, resulting in a plateauing of the signal. (vi) Once the initiation rate shuts off, no new RNAP molecules are added and the overall fluorescence signal starts to decrease due to cleavage of the nascent transcripts still on the gene. The MS2 and PP7 fluorescent signals are rescaled to be in the same arbitrary units with the calibration factor *α*. (Data in (B) adapted from ***Garcia et al. (2013)*** with the line representing the mean and error bars representing the standard error across 24 embryos.)

The steps of the transcription cycle can be coupled with each other. For example, elongation rates contribute to determining mRNA cleavage and RNA polymerase (RNAP) termination efficiency (***Pinto et al., 2011***; ***Hazelbaker et al., 2013***; ***Fong et al., 2015***; ***Liu et al., 2017***), and functional linkages have been demonstrated between transcription initiation and termination (***Moore and Proudfoot, 2009***; ***Mapendano et al., 2010***). Nonetheless, initiation, elongation, and transcript cleavage have largely been studied in isolation. In order to dissect the entire transcription cycle, it is necessary to develop a holistic approach that makes it possible to understand how the regulation of each step dictates mRNA production and to unearth potential couplings among these steps.

To date, the processes of the transcription cycle have mostly been studied in detail using *in vitro* approaches (***Bai et al., 2006***; ***Herbert et al., 2008***) or genome-wide measurements that require the fixation of cellular material and lack the spatiotemporal resolution to uncover how the regulation of the transcription cycle unfolds in real time (***Roeder, 1991***; ***Saunders et al., 2006***; ***Muse et al., 2007***; ***Core et al., 2008***; ***Fuda et al., 2009***; ***Churchman and Weissman, 2011***). Only recently has it become possible to dissect these processes in living cells and in their full dynamical complexity using tools such as MS2 or PP7 to fluorescently label nascent transcripts at single-cell resolution (***Bertrand et al., 1998***; ***Golding et al., 2005***; ***Chao et al., 2008***; ***Larson et al., 2011a***). These technological advances have yielded insights into, for example, intrinsic transcriptional noise in yeast (***Hocine et al., 2013***), kinetic splicing effects in human cells (***Coulon et al., 2014***), elongation rates in *Drosophila melanogaster* (***Garcia et al., 2013***; ***Fukaya et al., 2017***), and transcriptional bursting in mammalian cells (***Tantale et al., 2016***), *Dictyostelium* (***Chubb et al., 2006***; ***Muramoto et al., 2012***; ***Corrigan and Chubb, 2014***), fruit flies (***Garcia et al., 2013***; ***Lucas et al., 2013***; ***Bothma et al., 2014***; ***Fukaya et al., 2016***; ***Falo-Sanjuan et al., 2019***; ***Lammers et al., 2020***) and *Caenorhabditis elegans* (***Lee et al., 2019***).

Despite the great promise of MS2 and PP7, using these techniques to comprehensively analyze the transcription cycle is hindered by the fact that the signal from these *in vivo* RNA-labeling technologies convolves contributions from all aspects of the cycle. Specifically, the fluorescence signal from nascent RNA transcripts persists throughout the entire cycle of transcript initiation, elongation, and cleavage; further, a single gene can carry many tens of transcripts. Thus, at any given point, an MS2 or PP7 signal reports on the contributions of transcripts in various stages of the transcription cycle (***Ferraro et al., 2016***). Precisely interpreting an MS2 or PP7 signal therefore demands an integrated approach that accounts for this complexity.

Here, we present a method for analyzing live-imaging data from the MS2 and PP7 techniques in order to dynamically characterize the steps—initiation, elongation, and cleavage—of the full transcription cycle at single-cell resolution. While the transcription cycle is certainly more nuanced and can include additional effects such as sequence-dependent pausing (***Gaertner and Zeitlinger, 2014***), we view the quantification of these effective parameters as a key initial step for testing theoretical models. This method combines a dual-color MS2/PP7 fluorescent reporter (***Hocine et al., 2013***; ***Coulon et al., 2014***; ***Fukaya et al., 2017***) with Bayesian statistical inference techniques and quantitative modeling. As a proof of principle, we applied this analysis to the transcription cycle of a *hunchback* reporter gene in the developing embryo of the fruit fly *Drosophila melanogaster*. We validate our approach by comparing our inferred average initiation and elongation rates with previously reported results.

Crucially, our analysis also delivered novel single-cell statistics of the whole transcription cycle that were previously unmeasurable using genome-wide approaches, making it possible to generate distributions of parameter values necessary for investigations that go beyond simple population-averaged analyses (***Raj et al., 2006***; ***Zenklusen et al., 2008***; ***Sanchez et al., 2011***; ***Coulon et al., 2013***; ***Little et al., 2013***; ***Sanchez et al., 2013***; ***Sanchez and Golding, 2013***; ***Jones et al., 2014***; ***Serov et al., 2017***; ***Lucas et al., 2018***; ***Zoller et al., 2018***; ***Ali et al., 2020***; ***Filatova et al., 2020***). We show that, by taking advantage of time-resolved data, our inference is able to filter out uncorrelated noise, such as that originating from random measurement error, in these distributions and retain sources of correlated variability (such as biological and systematic noise. By combining these statistics with theoretical models, we revealed substantial variability in RNAP stepping rates between individual molecules, demonstrating the utility of our approach for testing hypotheses of the molecular mechanisms underlying the transcription cycle and its regulation.

This unified analysis enabled us to investigate couplings between the various transcription cycle parameters at the single-cell level, whereby we discovered a surprising correlation of cleavage rates with nascent transcript densities. These discoveries illustrate the potential of our method to sharpen hypotheses of the molecular processes underlying the regulation of the transcription cycle and to provide a framework for testing those hypotheses.

## Results

To quantitatively dissect the transcription cycle in its entirety from live imaging data, we developed a simple model (Fig. 1A) in which RNAP molecules are loaded at the promoter of a gene of total length *L* with a time-dependent loading rate *R*(*t*). For simplicity, we assume that each individual RNAP molecule behaves identically and independently: there are no interactions between molecules. While this assumption is a crude simplification, it nevertheless allows us to infer effective average transcription cycle parameters.

We parameterize this *R*(*t*) as the sum of a constant term ⟨*R*⟩ that represents the mean, or time-averaged, rate of initiation, and a small temporal fluctuation term given by δ*R*(*t*) such that *R*(*t*) = ⟨*R*⟩ + δ*R*(*t*). This mean-field parameterization is motivated by the fact that many genes are well approximated by constant rates of initiation (***Garcia et al., 2013***; ***Lucas et al., 2013***; ***Eck et al., 2020***; ***Lammers et al., 2020***). The fluctuation term δ*R*(*t*) allows for slight time-dependent deviations from the mean initiation rate. As a result, this term makes it possible to account for time-dependent behavior that can occur over the course of a cell cycle once the promoter has turned on. After initiation, each RNAP molecule traverses the gene at a constant, uniform elongation rate *v*_*elon*_. Upon reaching the end of the gene, there follows a deterministic cleavage time, *τ*_*cleave*_, after which the nascent transcript is cleaved.

We do not consider RNAP molecules that do not productively initiate transcription (***Darzacq et al., 2007***) or that are paused at the promoter (***Core et al., 2008***), as they will provide no experimental readout. Based on experimental evidence (***Garcia et al., 2013***), we assume that these RNAP molecules are processive, such that each molecule successfully completes transcription, with no loss of RNAP molecules before the end of the gene (see Section S5 for a validation of this hypothesis).

### Dual-color reporter for dissecting the transcription cycle

As a case study, we investigated the transcription cycle of early embryos of the fruit fly *D. melanogaster*. Specifically, we focused on the P2 minimal enhancer and promoter of the *hunchback* gene during the 14th nuclear cycle of development; the gene is transcribed in a step-like pattern along the anterior-posterior axis of the embryo with a 26-fold modulation in overall mRNA count between the anterior and posterior end (Fig. 1B; ***Driever and Nusslein-Volhard*** (***1989***); ***Margolis et al***. (***1995***); ***Perry et al. (2012***); ***Garcia et al. (2013***)). As a result, the fly embryo provides a natural modulation in mRNA production rates, with the position along the anterior-posterior axis serving as a proxy for mRNA output.

To visualize the transcription cycle, we utilized the MS2 and PP7 systems for live imaging of nascent RNA production (***Garcia et al., 2013***; ***Lucas et al., 2013***; ***Fukaya et al., 2016***). Using a two-color reporter construct similar to that reported in ***Hocine et al. (2013***), ***Coulon et al. (2014***), and ***Fukaya et al***. (***2017***), we placed the MS2 and PP7 stem loop sequences in the 5’ and 3’ ends, respectively, of a transgenic *hunchback* reporter gene (Fig. 1C; see Fig. S1 for more construct details). The *lacZ* sequence and a portion of the *lacY* sequence from *Escherichia coli* were placed as a neutral spacer (***Chen et al., 2012***) between the MS2 and PP7 stem loops.

As an individual RNAP molecule transcribes through a set of MS2/PP7 stem loops, constitutively expressed MCP-mCherry and PCP-GFP fusion proteins bind their respective stem loops, resulting in sites of nascent transcript formation that appear as fluorescent puncta under a laser-scanning confocal microscope (Fig. 2A and Video S1). The fluorescent signals did not exhibit noticeable photobleaching (Section S2 and Fig. S2). Since *hunchback* becomes transcriptionally active at the start of the nuclear cycle before slowly decaying into a transcriptionally silent state (***Garcia et al., 2013***; ***Liu et al., 2013***; ***Liu and Ma, 2015***), we restrict our analysis to the initial 18 minute window after mitosis where the promoter remains active.

**Figure 2.**
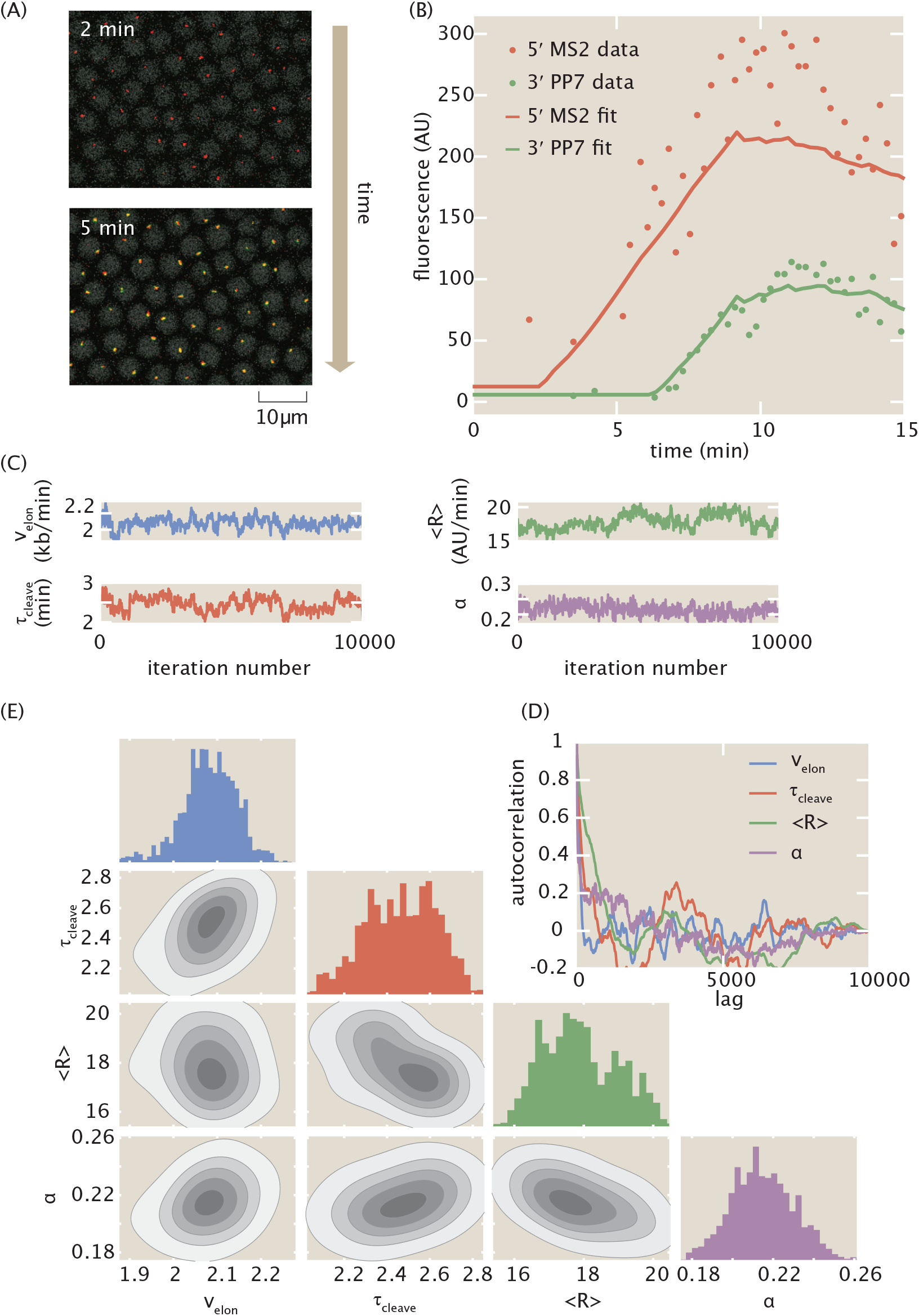
MCMC inference procedure. (A) Snapshots of confocal microscopy data over time, with MS2-mCherry (red) and PP7-eGFP (green) puncta reporting on transcription activity. Gray circles correspond to iRFP-labeled histones whose fluorescence is used as a fiduciary marker for cell nucleus segmentation (see Materials and Methods for details). (B) Sample single-cell MS2 and PP7 fluorescence (points) together with best-fits of the model using MCMC inference (curves). (C) Raw MCMC inference chains for the elongation rate *v*_*elon*_, cleavage time *τ*_*cleave*_, mean initiation rate ⟨*R*⟩, and calibration factor e*α* for the inference results of a sample single cell. (D) Auto-correlation function for the raw chains in (A) as a function of lag (i.e. inference sample number). (E) Corner plot of the raw chains shown in (C).

The intensity of the puncta in each color channel is linearly related to the number of actively transcribing RNAP molecules that have elongated past the location of the associated stem loop sequence (***Garcia et al., 2013***), albeit with different arbitrary fluorescence units. After reaching the end of the gene, which contains the 3’UTR of the e*α-tubulin* gene (***Chen et al., 2012***), the nascent RNA transcript undergoes cleavage. Because the characteristic timescale of mRNA diffusion is about two order of magnitudes faster than the time resolution of our experiment, we approximate the cleavage of a single transcript as resulting in the instantaneous loss of its associated fluorescent signal in both channels (Section S3). We included a few additional parameters in our model to make it compatible with this experimental data: a calibration factor e*α* between mCherry and eGFP intensities, a time of transcription onset *t*_*on*_ after mitosis at which the promoter switches on, and basal levels of fluorescence in each channel *MS*2_*basal*_ and *PP*7_*basal*_ (see Section S1 for more details). The qualitative relationship between the model parameters and the fluorescence data is described in Figure 1D, which considers the case of a pulse of constant initiation rate.

### Transcription cycle parameter inference using Markov Chain Monte Carlo

We developed a statistical framework to estimate transcription-cycle parameters (Fig. 1A) from fluorescence signals. Time traces of mCherry and eGFP fluorescence intensity are extracted from microscopy data such as shown in Figure 2A and Video S1 to produce a dual-signal readout of nascent RNA transcription at single-cell resolution (Fig. 2B, data points; see Methods and Materials for details). To extract quantitative insights from the observed fluorescence data, we used the established Bayesian inference technique of Markov Chain Monte Carlo (MCMC) (***Geyer, 1992***) to infer the effective parameter values in our simple model of transcription: the calibration factor between mCherry and eGFP intensities *α*, the time-dependent transcription initiation rate, separated into the constant term ⟨*R*⟩ and fluctuations δ*R*(*t*), the elongation rate *v*_*elon*_, the cleavage time *τ*_*cleave*_, the time of transcription onset *t*_*on*_, and the basal levels of fluorescence in each channel *MS*2_*basal*_ and *PP*7_*basal*_.

The details of the inference procedure are described in Section S4.1. Briefly, the inference was run separately for each single cell, yielding chains of sampled parameter values (Fig. 2C). These resulting chains exhibited rapid mixing and rapidly decaying auto-correlation functions (Fig. 2D), indicative of reliable fits. Corner plots of the fits indicated reasonable posterior distributions (Fig. 2E).

From these single-cell fits, the mean value of each parameter’s chain was retained for further analysis. The final dataset was produced by filtering with an automated procedure that relied on overall fit quality (Section S4.3 and Fig. S4). This curation procedure did not introduce noticeable bias in the results (Fig. S4G-I). A small minority of the rejected cells (Fig. S4E) exhibited highly time-dependent behavior reminiscent of transcriptional bursting (***Rodriguez and Larson, 2020***), which lies outside the scope of our model and is explored more in the Discussion. A sample fit is shown in Figure 2B. To aggregate the results, we constructed a distribution from the inferred parameter from each single-cell. Intra-embryo variability between single cells was greater than inter-embryo variability (Section S6 and Fig. S6). As a result, unless stated otherwise, all statistics reported here were aggregated across 355 single cells combined between 7 embryos, and all shaded errors reflect the standard error of the mean.

### MCMC successfully infers calibration between eGFP and mCherry intensities

Due to the fact that the MS2 and PP7 stem loop sequences were associated with mCherry and eGFP fluorescent proteins, respectively, the two experimental fluorescent signals possessed different arbitrary fluorescent units, related by the scaling factor *α* given by

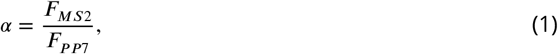

where *F*_*MS*2_ and *F*_*PP*7_ are the fluorescence values generated by a fully transcribed set of MS2 and PP7 stem loops, respectively. Although *α* has units of *AU*_MS2_/*AU*_PP7_, we will express e*α* without units in the interest of clarity of notation.

We inferred single-cell values of *α* using the inference methodology. As shown in the blue histogram in Figure 3A, our inferred values of e*α* possessed a mean of 0.145 ± 0.004 (SEM) and a standard deviation of 0.068.

**Figure 3.**
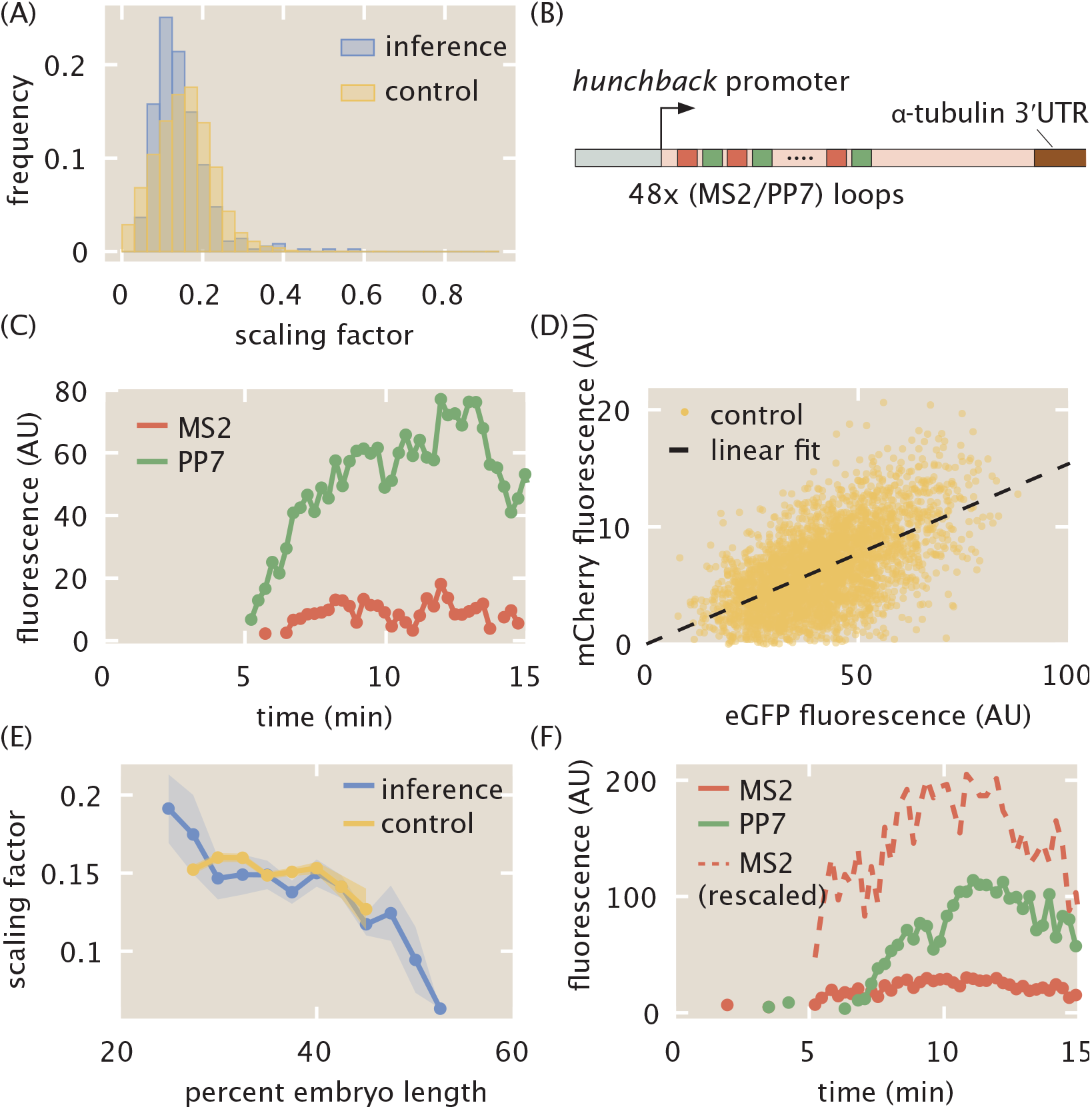
Calibration of MS2 and PP7 fluorescence signals. (A) Histogram of inferred values of e*α* at the single-cell level from inference (blue), along with histogram of e*α* values from the control experiment (yellow). (B) Schematic of construct used to measure the calibration factor e*α* using 24 interlaced MS2/PP7 loops each (48 loops in total). (C) Sample single-cell MS2 (red) and PP7 (green) traces from this control experiment. (D) Scatter plot of MS2 and PP7 fluorescence values for each time point (yellow) along with linear best fit (black) resulting in e*α* = 0.154 ± 0.001. (E) Position-dependent mean value of e*α* in both the inference (blue) and the control experiment (yellow). (F) Representative raw and rescaled MS2 and PP7 traces for a sample single cell in the inference data set. (A,D,E, data were collected for 314 cells across 4 embryos for the interlaced reporter, and for 355 cells across 7 embryos for the reporter with MS2 on the 5’ and PP7 on the 3’ of the gene (Fig. 1C); shaded regions in (E) reflect standard error of the mean. Measurement conditions for both experiments are described in Materials and Methods.)

As an independent validation, we measured *α* by using another two-color reporter, consisting of 24 alternating, rather than sequential, MS2 and PP7 loops (***Wu et al., 2014***; ***Chen et al., 2018***; ***Child et al., 2020***) inserted at the 5’ end of our reporter construct (Fig. 3B). Thus, this reporter had a total of 48 stem loops, with 24 each of MS2 and PP7.

Figure 3C shows a representative trace of a single spot containing our calibration construct (see Video S2 for full movie). For each time point, the mCherry fluorescence in all measured single-cell traces was plotted against the corresponding eGFP fluorescence (Fig. 3D, yellow points). The mean e*α* was then calculated by fitting the resulting scatter plot to a line going through the origin (Fig. 3D, black line). The best-fit slope yielded the experimentally calculated value of e*α* = 0.154 ± 0.001 (SEM). A distribution for e*α* was also constructed by dividing the mCherry fluorescence by the corresponding eGFP fluorescence for each datapoint in Figure 3D, yielding the histogram in Figure 3A (yellow), which possessed a standard deviation of 0.073. Our independent calibration agreed with our inference, thus validating the infererred values of e*α*.

Interestingly, binning the cells by position along the embryo revealed a slight position dependence in the scaling factor. As shown in Figure 3E, both the directly measured and inferred e*α* displayed higher values in the anterior, about 0.15, and lower values in the posterior, about 0.1. The fact that this position dependence is observed in both in the calibration experiments and inference suggests that this spatial modulation in the value of e*α* is not an artifact of the constructs or our analysis, but a real feature of the system. We speculate that this spatial dependence could stem from differential availability of MCP-mCherry and PCP-GFP along the embryo, leading to a modulation in the maximum occupancy of the MS2 stem loops versus the PP7 stem loops (***Wu et al., 2012***).

Regardless, our data demonstrate that the inferred and calibrated e*α* can be used interchangeably, obviating the need for the control. Thus, the MS2 signals for each single cell could be rescaled to the same units as the PP7 signal (Fig. 3F) within a single experiment, greatly increasing the power of the inference methodology. All plots in the main text and supplementary information, unless otherwise stated, reflect these rescaled values using the overall mean value of e*α* = 0.145 obtained from the inference.

### Inference of single-cell initiation rates recapitulates and improves on previous measurements

After validating the accuracy of our inference method in inferring transcription initiation, elongation, and cleavage dynamics using simulated data (Section S4.4 and Fig. S5), we inferred these transcriptional parameters for the *hunchback* reporter gene as a function of the position along the anterior-posterior axis of the embryo. The suite of quantitative measurements on the transcription cycle produced by the aggregated inference results is shown in Figures 4A, C, E, and F. Full distributions of these parameters can be found in Fig. S7.

**Figure 4.**
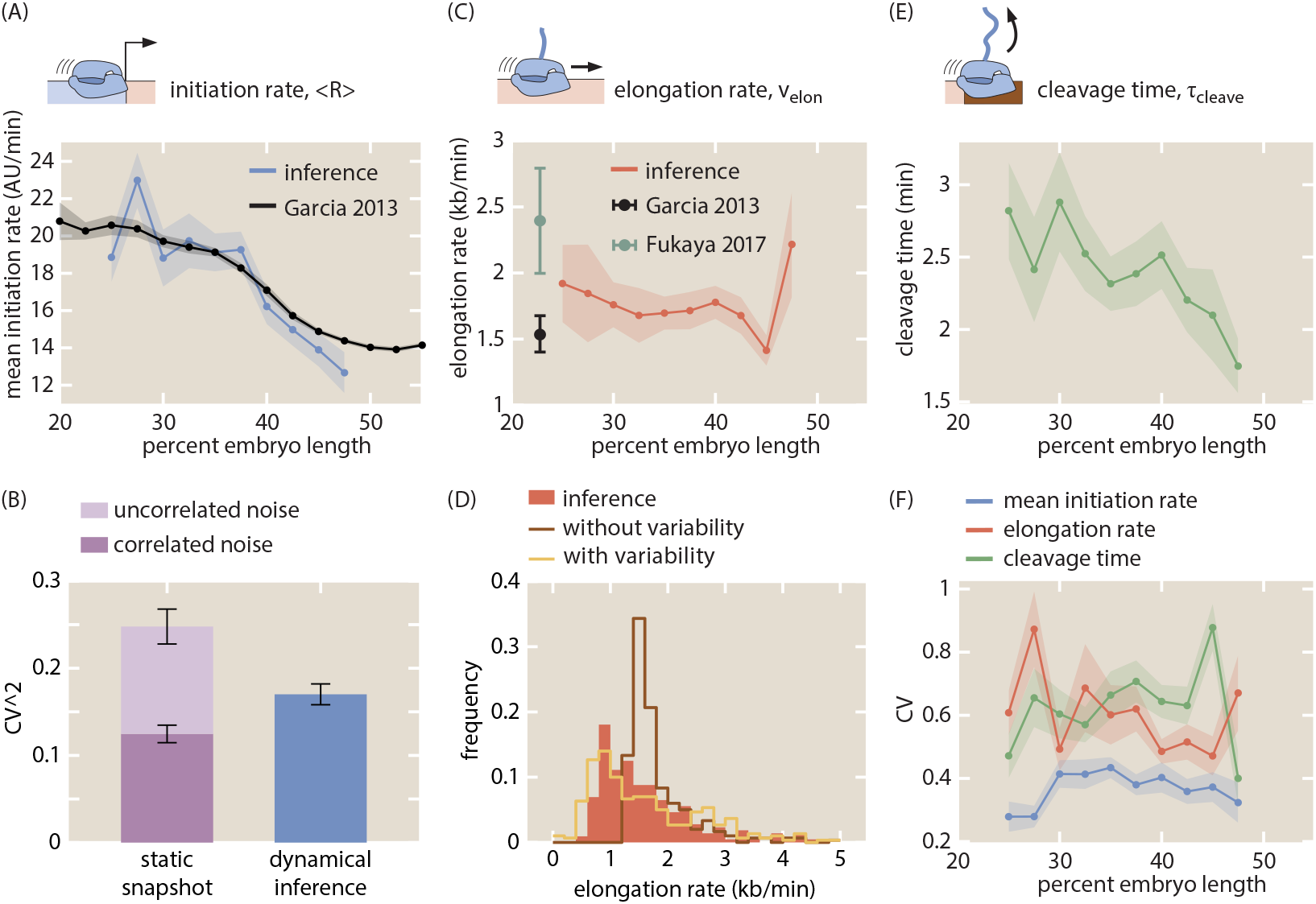
Inferred transcription-cycle parameters. (A) Mean inferred transcription initiation rate as a function of embryo position (blue), along with rescaled previously reported results (black, ***Garcia et al***. (***2013***)). (B) Comparison of the squared CV of the mean initiation rate inferred using our approach (blue) or obtained from examining the fluorescence of transcription spots in a single snapshot (light plus dark purple). While snapshots captured a significant amount of uncorrelated noise (light purple), our inference accounts mostly for correlated noise (compare blue and dark purple). See Section S8 and Fig. S8 for details. (C) Inferred elongation rate as a function of embryo position (red), along with previously reported results (black, ***Garcia et al***. (***2013***); teal, ***Fukaya et al***. (***2017***)). (D) Distribution of inferred single-cell elongation rates in the anterior 40% of embryo (red), along with best fit to mean and standard deviation using single-molecule simulations with and without RNAP-to-RNAP variability (gold and brown, respectively, see Section S10 for details). (E) Inferred cleavage time as a function of embryo position. (F) CV of the mean initiation rate (blue), elongation rate (red), and cleavage time (green) as a function of embryo position. (A, C, E, shaded error reflects standard error of the mean across 355 nuclei in 7 embryos, or of previously reported mean results; B, F, shaded error or black error bars represent bootstrapped standard errors of the CV or CV^2^ for 100 bootstrap samples each; C, error bars reflect standard error of the mean for ***Garcia et al***. (***2013***) and lower (25%) and upper (75%) quintiles of the full distribution from ***Fukaya et al***. (***2017***).)

Control of initiation rates is one of the predominant, and as a result most well-studied, strategies for gene regulation (***Roeder, 1991***; ***Spitz and Furlong, 2012***; ***Lenstra et al., 2016***). Thus, comparing our inferred initiation rates with previously established results comprised a crucial benchmark for our methodology. Our inferred values of the mean initiation rate ⟨*R*⟩ exhibited a step-like pattern along the anterior-posterior axis of the embryo, qualitatively reproducing the known *hunchback* expression profile (Fig. 4A, blue). As a point of comparison, we also examined the mean initiation rate measured by ***Garcia et al***. (***2013***), which was obtained by manually fitting a trapezoid (Figure 1D) to the average MS2 signal (Fig. 4A, black). The quantitative agreement between these two dissimilar analysis methodologies demonstrates that our inference method can reliably extract the average rate of transcription initiation across cells.

Measurements of cell-to-cell variability in transcription initiation rate have uncovered, for example, the existence of transcriptional bursting and mechanisms underlying the establishment of precise developmental boundaries (***Raj et al., 2006***; ***Sanchez and Golding, 2013***; ***Zenklusen et al., 2008***; ***Little et al., 2013***; ***Jones et al., 2014***; ***Lucas et al., 2018***; ***Zoller et al., 2018***). Yet, to date, these studies have mostly employed techniques such as single-molecule FISH to count the number of nascent transcripts on a gene or the number of cytoplasmic mRNA molecules (***Femino et al., 1998***; ***Raj et al., 2006***; ***Pare et al., 2009***; ***Zenklusen et al., 2008***; ***So et al., 2011***; ***Boettiger and Levine, 2013***; ***Little et al., 2013***; ***Jones et al., 2014***; ***Senecal et al., 2014***; ***Padovan-Merhar et al., 2015***; ***Xu et al., 2015***; ***Skinner et al., 2016***; ***Bartman et al., 2016***; ***Hendy et al., 2017***; ***Zoller et al., 2018***). In principle, these techniques do not report on the variability in transcription initiation alone; they convolve this measurement with variability in other steps of the transcription cycle (***Padovan-Merhar et al., 2015***; ***Lenstra et al., 2016***).

Our inference approach isolates the transcription initiation rate from the remaining steps of the transcription cycle at the single-cell level, making it possible to calculate, for example, the coefficient of variation (CV; standard deviation divided by the mean) of the mean rate of initiation. Our results yielded values for the CV along the embryo that were fairly uniform, with a maximum value of around 40% (Fig. 4F, blue). This value is roughly comparable to that obtained for *hunchback* using single-molecule FISH (***Little et al., 2013***; ***Xu et al., 2015***; ***Zoller et al., 2018***).

One of the challenges in measuring CV values, however, is that informative biological variability is often convolved with undesired experimental noise, such as experimental measurement noise inherent to fluorescence microscopy. In general, this experimental noise can contain both random, uncorrelated components as well as systematic components, the latter of which combines with actual biological variability to form overall correlated noise. Although we currently cannot entirely separate biological variability from experimental noise with our data and inference method, a strategy for at least separating uncorrelated from correlated was recently implemented in the context of snapshot-based fluorescent data (***Zoller et al., 2018***). By utilizing a dual-color measurement of the same biological signal, one can separate the total variability in a dataset into uncorrelated measurement noise and correlated noise, which includes components such as true biological variability and systematic measurement error.

Building on this strategy, we first took a single snapshot from our live-imaging data and calculated the total squared CV of the fluorescence of spots at a single time point (Fig. 4B, dark plus light purple). Compared to the squared CV from the inferred mean initiation rate (Fig. 4B, blue), the squared CV from the snapshot was larger by about 0.1, suggesting that the inference method reported on a somewhat lower level of overall variability.

To investigate this disparity in measured variability further, we then rewrote the squared CV from the snapshot approach as the sum of uncorrelated and correlated noise components

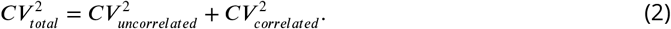

The magnitudes of each noise component were estimated by using the data from the interlaced reporter introduced in Figure 3B. To do so, we utilized the fact that, in principle, the mCherry and GFP signals from this experiment reflected the same underlying biological process, and assumed that deviations between the two signals were a result of uncorrelated measurement noise. Thus, we could apply the two-color formalism introduced in ***Elowitz et al***. (***2002***) to calculate the uncorrelated and correlated noise components from snapshots taken from the interlaced reporter construct (see Section S8 and Figure S8 for more details).

The bar graph shown in Figure 4B shows that, once the uncorrelated noise (light purple) is subtracted from the total noise of our snapshot-based measurement, the remaining correlated variability (dark purple), which includes the biological variability, is slightly lower than the variability of our inference results (blue). Thus, our inference mostly captures correlated variability and filters out the bulk of the uncorrelated noise, similarly to techniques such as single-molecule FISH (***Zoller et al., 2018***) but with the added advantage of also being able to resolve temporal information. Because such fixed tissue techniques ultimately provide static measurements that convolve signals from transcription initiation with those of elongation and cleavage, it is important to note that this is a qualitative comparison between the ability of fixed-tissue and live-imaging to separate correlated and uncorrelated variability. Thus, our results further validate our approach and demonstrate its capability to capture measures of cell-to-cell variability in the transcription cycle with high precision.

### Elongation rate inference reveals single-molecule variability in RNAP stepping rates

Next, we investigated the ability of our inference approach to report on the elongation rate *v*_*elon*_. Nascent RNA elongation plays a prominent role in gene regulation, for example, in dosage compensation in *Drosophila* embryos (***Larschan et al., 2011***), alternative splicing in human cells (***De La Mata et al., 2003***; ***Batsché et al., 2006***), and gene expression in plants (***Wu et al., 2016***). Our method inferred an elongation rate *v*_*elon*_ that was relatively constant along the embryo (Fig. 4C), lending support to previous reports indicating a lack of regulatory control of the elongation rate in the early fly embryo (***Fukaya et al., 2017***). We measured a mean elongation rate of 1.72 ± 0.05 kb/min (SEM; n = 355), consistent with previous measurements of the fly embryo (Fig. 4C, black and teal; ***Garcia et al***. (***2013***); ***Fukaya et al***. (***2017***)), as well as with measurements from other techniques and model organisms, which range from about 1 kb/min to upwards of 4 kb/min (***Golding et al., 2005***; ***Darzacq et al., 2007***; ***Boireau et al., 2007***; ***Ardehali and Lis, 2009***; ***Hocine et al., 2013***; ***Coulon et al., 2014***; ***Fuchs et al., 2014***; ***Tantale et al., 2016***; ***Lenstra et al., 2016***). In addition, the CV of the elongation rate was roughly uniform across embryo position (Fig. 4F, red).

Like cell-to-cell variability in transcription initiation, single-cell distributions of elongation rates can provide crucial insights into, for example, promoter-proximal pausing (***Serov et al., 2017***), traffic jams (***Klumpp and Hwa, 2008***; ***Klumpp, 2011***), transcriptional bursting (***Choubey et al., 2015, 2018***), and noise propagation (***Ali et al., 2020***). While genome-wide approaches have had huge success in measuring mean properties of elongation (***Core et al., 2008***; ***Carrillo Oesterreich et al., 2010***), they remain unable to resolve single-cell distributions of elongation rates. We examined the statistics of single-cell elongation rates in the anterior 40% of the embryo, where the initiation rate was roughly constant, and inferred a broad distribution of elongation rates with a standard deviation of around 1 kb/min and a long tail extending to values upwards of 4 kb/min (Fig. 4D, red). This large spread was consistent with observations of large cell-cell variability in elongation rates (***Palangat and Larson, 2012***; ***Lenstra et al., 2016***) using a wide range of techniques, as well as with measurements from similar two-color live imaging experiments (***Hocine et al***. (***2013***); ***Fukaya et al***. (***2017***); Section S9; Fig. S9).

To illustrate the resolving power of examining elongation rate distributions, we performed theoretical investigations of cell-to-cell variability in this transcription cycle parameter. Following ***Klumpp and Hwa*** (***2008***), we considered a model where RNAP molecules stochastically step along a gene and cannot overlap or pass each other (Section S10). The model simulated MS2 and PP7 fluorescences that were then run through the inference procedure, in order to account for the presence of inferential noise (Section S4.4).

First, we considered a scenario where the stepping rate of each RNAP molecule is identical. In this case, the sole driver of cell-to-cell variability is the combination of stochastic stepping behavior with traffic jamming due to steric hindrance of RNAP molecules. As shown in brown in Figure 4D, this model cannot account for the wide distribution of observed single-cell elongation rates.

In contrast, by allowing for substantial variability in the elongation rate of individual RNAP molecules, the model can reproduce the empirical distribution of single-cell elongation rates. As shown in gold in Figure 4D, the model can quantitatively approximate the inferred distribution within error (Fig. S10D). This single-molecule variability is consistent with *in vitro* observations of substantial molecule-to-molecule variability in RNAP elongation rates (***Tolić-Nørrelykke et al., 2004***; ***Larson et al., 2011b***), thus demonstrating the ability of our approach to engage in the *in vivo* dissection of the transcription cycle at the single-molecule level.

### Inference reveals functional dependencies of cleavage times

Finally, we inferred values of the cleavage time *τ*_*cleave*_. Through processes such as alternative polyadenylation (***Tian and Manley, 2016***; ***Jung et al., 2009***) and promoter-terminator crosstalk (***Moore and Proudfoot, 2009***; ***Mapendano et al., 2010***), events at the 3’ end of a gene exert substantial influence over overall transcription levels (***Bentley, 2014***). Although many investigations of mRNA cleavage and RNAP termination have been carried out in fixed-tissue samples (***Richard and Manley, 2009***; ***Kuehner et al., 2011***), live-imaging studies with single-cell resolution of this important process remain sparse; some successes have been achieved in yeast and in mammalian cells (***Lenstra et al., 2016***). We inferred a mean mRNA cleavage time in the range of 1.5-3 min (Fig. 4E), consistent with values obtained from live imaging in yeast (***Larson et al., 2011a***) and mammalian cells (***Boireau et al., 2007***; ***Darzacq et al., 2007***; ***Coulon et al., 2014***; ***Tantale et al., 2016***). Interestingly, as shown in Figure 4E, the inferred mRNA cleavage time was dependent on anterior-posterior positioning along the embryo, with high values (∼3 min) in the anterior end and lower values toward the posterior end (∼1.5 min). While the reasons for this position dependence are unknown, such dependence could result from the presence of a spatial gradient of a molecular species that regulates cleavage. Importantly, such a modulation could not have been easily revealed using genome-wide approaches that, by necessity, average information across multiple cells.

The CV of the cleavage time slightly increased toward the posterior end of the embryo (Fig. 4F, green). Thus, although cleavage remains an understudied process compared to initiation and elongation, both theoretically and experimentally, these results provide the quantitative precision necessary to carry out such mechanistic analyses.

### Uncovering single-cell mechanistic correlations between transcription cycle parameters

In addition to revealing trends in average quantities of the transcription cycle along the length of the embryo, the simultaneous nature of the inference afforded us the unprecedented ability to investigate single-cell correlations between transcription-cycle parameters. We used the Spearman rank correlation coefficient (ρ) as a non-parametric measure of inter-parameter correlations. The mean initiation rate and the cleavage time exhibited a negative correlation (ρ = −0.52, *p* ≈ 0; Fig. 5A). This negative correlation at the single-cell level should be contrasted with the positive relation between these magnitudes at the position-averaged level, where the mean initiation rate and cleavage time both increased in the anterior of the embryo (Fig. 4A and E). Thus, our analysis unearthed a quantitative relationship that was obscured by a naive investigation of spatially averaged quantities, an approach often used in fixed (***Zoller et al., 2018***) and live-imaging (***Lammers et al., 2020***) studies, as well as in genome-wide investigations (***Combs and Eisen, 2017***; ***Haines and Eisen, 2018***). We also detected a small negative correlation (ρ = −0.21, *p* = 5 × 10^−5^) between elongation rates and mean initiation rates (Fig. 5B). Finally, we detected a small positive correlation (ρ = 0.35, *p* = 2 × 10^−11^) between cleavage times and elongation rates (Fig. 5C). These results are consistent with prior studies implicating elongation rates in 3’ processes such as splicing and alternative polyadenylation: slower elongation rates increased cleavage efficiency (***De La Mata et al., 2003***; ***Pinto et al., 2011***).

**Figure 5.**
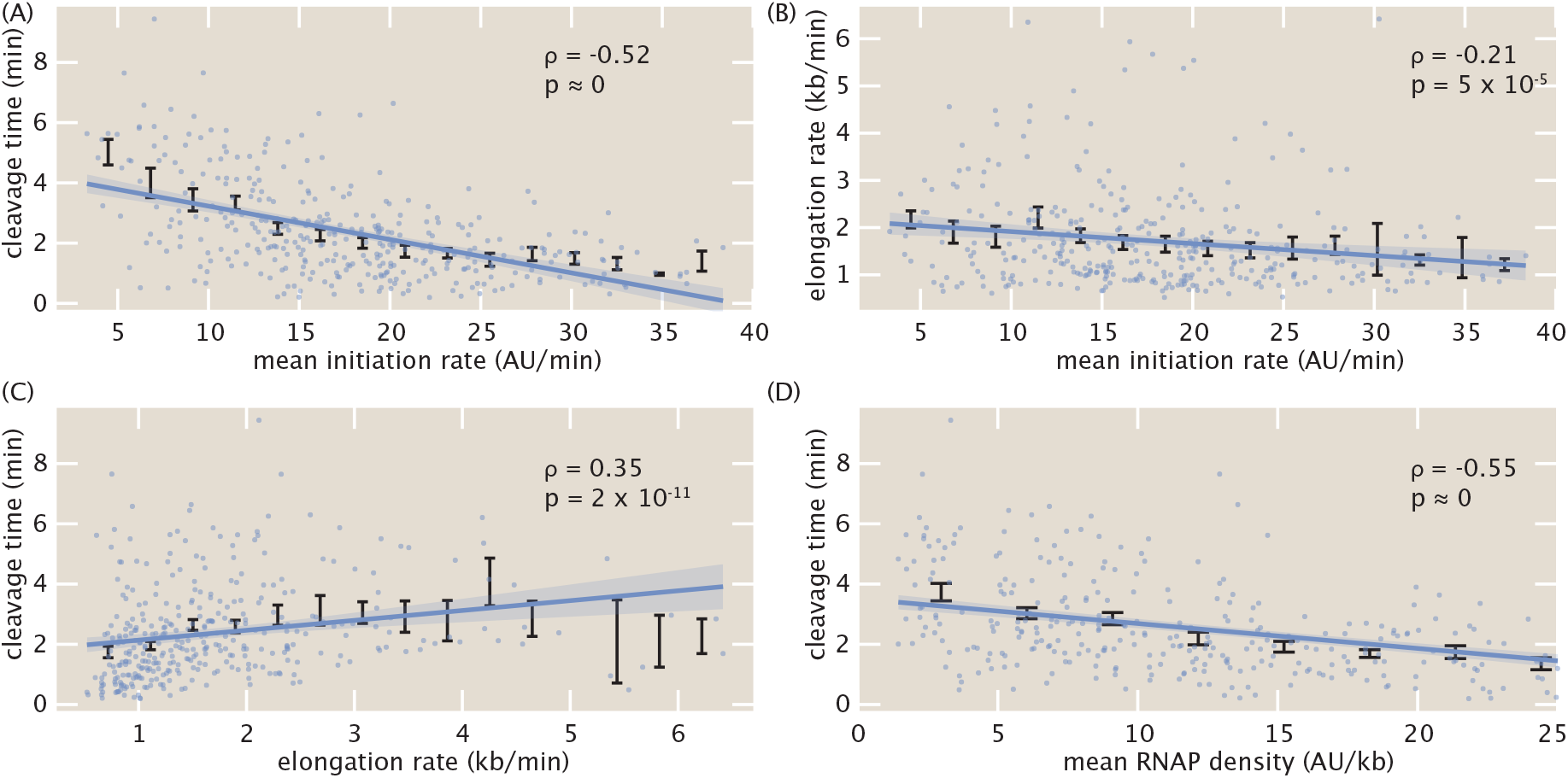
Single-cell correlations between transcription cycle parameters. Spearman rank correlation coefficients and associated p-values between (A) mean initiation rate and cleavage time, (B) mean initiation rate and elongation rate, (C) elongation rate and cleavage time, and (D) mean RNAP density and cleavage time. Blue points indicate single-cell values; black points and error bars indicate mean and SEM, respectively, binned across x-axis values. Lines and shaded regions indicate generalized linear model fit and 95% confidence interval, respectively, and are shown for ease of visualization (see Materials and Methods for details).

The observed negative correlation between cleavage time and mean initiation rate (Fig. 5A), in conjunction with the positive correlation between cleavage time and elongation rate (Fig. 5C), suggested a potential underlying biophysical control parameter: the mean nascent transcript density on the reporter gene body ρ given by

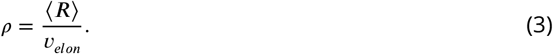

Possessing units of (AU/kb), this mean transcript density estimates the average number of nascent RNA transcripts per kilobase of template DNA. Plotting the cleavage time as a function of the mean transcript density yielded a negative correlation (ρ = −0.55, *p* ≈ 0) that was stronger than any of the other correlations between transcription-cycle parameters at the single-cell level (Fig. 5D). Mechanistically, the correlation between cleavage time and mean transcript density suggests that, on average, more closely packed nascent transcripts at the 3’ end of a gene cleave faster.

Further investigations using simulations indicated that this relationship did not arise from spurious correlations in the inference procedure itself (Section S4.4 and Fig. S5E-H), but rather captured real correlations in the data. Furthermore, although the four inter-parameter correlations investigated here only used mean values obtained from the inference methodology, a Monte Carlo simulation involving the full Bayesian posterior distribution confirmed the significance of the results (Section S11 and Fig. S11).

Using an absolute calibration for a similar reporter gene (***Garcia et al., 2013***) led to a rough scaling of 1 AU ≈ 1 molecule corresponding to a maximal RNAP density of about 20 RNAP molecules/kb in Figure 5D. With a DNA footprint of 40 bases per molecule (***Selby et al., 1997***), this calculation suggests that, in this regime, RNAP molecules are densely distributed, occupying about 80% of the reporter gene. We hypothesize that increased RNAP density could lead to increased pausing as a result of traffic jams (***Klumpp and Hwa, 2008***; ***Klumpp, 2011***). Due to this pausing, transcripts would be more available for cleavage, increasing overall cleavage efficiency. Regardless of the particular molecular mechanisms underlying our observations, we anticipate that this ability to resolve single-cell correlations between transcription parameters, combined with perturbative experiments, will provide ample future opportunities for studying the underlying biophysical mechanisms linking transcription processes.

## Discussion

Over the last two decades, the genetically encoded MS2 (***Bertrand et al., 1998***) and PP7 (***Chao et al., 2008***) RNA labeling technologies have made it possible to measure nascent and cytoplasmic RNA dynamics *in vivo* in many contexts (***Golding et al., 2005***; ***Chubb et al., 2006***; ***Darzacq et al., 2007***; ***Larson et al., 2011a***; ***Garcia et al., 2013***; ***Lucas et al., 2013***; ***Hocine et al., 2013***; ***Coulon et al., 2014***; ***Bothma et al., 2014***; ***Lenstra et al., 2016***; ***Fukaya et al., 2016***; ***Tantale et al., 2016***; ***Fukaya et al., 2017***; ***Chen et al., 2018***; ***Dufourt et al., 2018***; ***Fritzsch et al., 2018***; ***Falo-Sanjuan et al., 2019***; ***Li et al., 2019***; ***Lee et al., 2019***; ***Lammers et al., 2020***; ***Eck et al., 2020***). However, such promising experimental techniques can only be as powerful as their underlying data-analysis infrastructure. For example, while initial studies using MS2 set the technological foundation for revealing transcriptional bursts in bacteria (***Golding et al., 2005***), single-celled eukaryotes (***Chubb et al., 2006***; ***Larson et al., 2009***), and animals (***Garcia et al., 2013***; ***Lucas et al., 2013***), only recently did analysis techniques become available to reliably obtain parameters such as transcriptional burst frequency, duration, and amplitude (***Coulon et al., 2014***; ***Desponds et al., 2016***; ***Corrigan et al., 2016***; ***Lammers et al., 2020***; ***Bowles et al., 2020***).

In this work, we established a novel method for inferring quantitative parameters of the entire transcription cycle—initiation, elongation and cleavage—from live imaging data of nascent RNA dynamics. Notably, this method offers high spatiotemporal resolution at the single-cell level, resolving aspects of transcriptional activity within the body of an organism and at sub-minute resolution. Furthermore, while our experimental setup utilized two fluorophores, we found that the calibration between their intensities could be inferred directly from the data (Fig. 3), rendering independent calibration and control experiments unnecessary.

After validating previously discovered spatial modulations in the mean initiation rate, we discovered an unreported modulation of the cleavage time with respect to embryo position that mirrored that of the mean initiation rate (Fig. 4E). Although such a relationship at first would suggest a positive correlation between initiation and cleavage, the presence of significant negative correlation at the single-cell level refutes this idea (Fig. 5A). Instead, we speculate that the spatial modulation of the cleavage time could instead underlie a coupling with a spatial gradient of some molecular factor that controls this transcription cycle parameter (***El Kaderi et al., 2009***), possibly due to effects such as gene looping (***O’Sullivan et al., 2004***; ***Tan-Wong et al., 2008***).

These features are unattainable by widespread, but still powerful, genome-wide techniques that examine fixed samples, such as global run-on sequencing (GRO-seq) to measure elongation rates *in vivo* (***Danko et al., 2013***; ***Jonkers and Lis, 2015***). Additionally, while fixed-tissue technologies such as single-molecule RNA-FISH provide superior spatial and molecular resolution to current live imaging technologies (***Little et al., 2013***; ***Zoller et al., 2018***), the fixation process necessarily prevents any sort of temporal single-cell analysis of the highly dynamic transcriptional processes.

### Dissecting the transcription cycle at the single-cell level

From elucidating the nature of mutations (***Luria and Delbruck, 1943***) and revealing mechanisms of transcription initiation (***Zenklusen et al., 2008***; ***Sanchez et al., 2011***; ***So et al., 2011***; ***Sanchez et al., 2013***; ***Sanchez and Golding, 2013***; ***Little et al., 2013***; ***Hocine et al., 2013***; ***Jones et al., 2014***; ***Xu et al., 2015***; ***Choubey et al., 2015***; ***Zoller et al., 2018***; ***Choubey et al., 2018***; ***Filatova et al., 2020***), transcription elongation (***Boettiger et al., 2011***; ***Serov et al., 2017***; ***Ali et al., 2020***), and translational control (***Cai et al., 2006***), to enabling the calibration of fluorescent proteins in absolute units (***Rosenfeld et al., 2005, 2006***; ***Teng et al., 2010***; ***Brewster et al., 2014***; ***Kim et al., 2016***; ***Bakker and Swain, 2019***), examining single-cell distributions through the lens of theoretical models has made it possible to extract molecular insights about biological function that are inaccessible through the examination of averaged quantities. The single-cell measurements afforded by our approach made it possible to infer full distributions of transcription parameters (Fig. 4B, D, and F). This single-cell resolution motivates a dialogue between theory and experiment for studying transcription initiation, elongation, and cleavage at the single-cell level.

We showed how our inferred distributions of initiation rates effectively filter out most uncorrelated measurement noise, which we expect to be dominated by experimental noise, while retaining information on sources of correlated noise, including underlying biological variability (Fig. 4B). Additionally, our theoretical model of elongation rate distributions make it possible to test mechanistic models of RNAP transit along the gene. While still preliminary and far from conclusive, our results suggest that cell-to-cell variability in elongation rates arises from single-molecule variability in stepping rates, and that processes such as stochasticity in stepping behavior and traffic jamming due to steric hindrance alone cannot account for the observed elongation rate distributions (Fig. 4D). Such statistics could then be harnessed to make predictions for future perturbative experiments that utilize, for example, mutated RNAP molecules with altered elongation rates (***Chen et al., 1996***) or reporter genes with differing spacer lengths between MS2 and PP7 stem loops sequences.

Finally, the simultaneous single-cell inference of transcription-cycle parameters granted us the novel capability to investigate couplings between transcription initiation, elongation, and cleavage, paving the way for future studies of mechanistic linkages between these processes. In particular, the observed coupling of the mRNA cleavage time with RNAP density (Fig. 5D) suggests future experiments utilizing, for example, orthogonal stem loops on either side of the 3’UTR as potential avenues for investigating mechanisms such as RNAP traffic jams (***Klumpp and Hwa, 2008***; ***Klumpp, 2011***), inefficient or rate-limiting nascent RNA cleavage (***Fong et al., 2015***; ***Jung et al., 2009***), and promoter-terminator looping (***Hampsey et al., 2011***). Other potential experiments could include perturbative effects, such as introducing inhibitors of transcription initiation, elongation, and/or cleavage and assessing the downstream impact on the inferred transcriptional parameters to see if the perturbed effects are separable or convolved between parameters.

### Comparison to existing analysis techniques

Our method provides a much-needed framework for applying statistical inference for the analysis of live imaging data of nascent transcription, expanding the existing repertoire of model-driven statistical techniques to analyze single-cell protein reporter data (***Heron et al., 2007***; ***Finkenstädt et al., 2008***; ***Suter et al., 2011***; ***Zechner et al., 2014***). In particular, compared to auto-correlation analysis of transcriptional signals (***Coulon and Larson, 2016***), another powerful method of analyzing live imaging transcription data, our method is quite complementary.

First, auto-correlation analysis typically requires a time-homogeneous transcript initiation process (***Coulon and Larson, 2016***), and benefits immensely from having experimental data acquired over long time windows to enhance the auto-correlation signal (although recent work has improved on the ability to analyze short time windows (***Desponds et al., 2016***)). In contrast, our model-driven inference approach can account for slight time dependence and can fit short time traces. This is of particular relevance to the fly embryo, where each cell cycle in early development is incredibly short (here, we only examined 18 minutes of data) and transcription initiation switches from OFF to ON and back to OFF within that timeframe.

Second, auto-correlation analysis depends strongly on signal-noise ratio, namely the ability to resolve single-or-few-transcript fluctuations in the number of actively transcribing polymerases on a gene (***Larson et al., 2011a***; ***Coulon et al., 2014***). Our approach, however, can be applied even if the signal-noise ratio can only resolve differences in transcript number of several transcripts, rather than just one.

Third, our model-driven approach benefits from explicitly parameterizing the various steps of the transcription cycle, allowing for the separation of processes such as elongation and cleavage. In contrast, while the auto-correlation technique has the advantage of not relying on a particular specific model, it does rely on unknown parameters such as the overall transcript dwell time, which is a combination of elongation and cleavage. Thus, it becomes harder to separate contributions from these different processes. Additionally, auto-correlation approaches cannot produce absolute rates of transcriptional processes, such as the quantified rates of mean transcription initiation obtained in this work.

### Future improvements

Future improvements to experimental or inferential resolution could sharpen precision of single-cell results, increasing confidence in the distributions obtained through this methodology. For example, technologies such as lattice light-sheet microscopy (***Chen et al., 2014***; ***Mir et al., 2017, 2018***) would vastly improve spatiotemporal imaging resolution and reduce uncertainty in measurements. While this increased resolution is unlikely to dramatically change the statistics reported here, it could potentially push the analysis regime to the single-molecule level, necessitating the parallel development of increasingly refined models that can account for stochasticity and fluctuations that are not resolved with bulk measurements. In addition, while our analysis restricted itself to consider only nascent RNA labeling technologies, this methodology could be extended to also examine mature labeled RNA in the nucleus and cytoplasm of an organism, providing a more complete picture of transcription.

One important caveat of our method is the failure to account for genes that undergo transcriptional bursting (***Rodriguez and Larson, 2020***). Here, the initiation rate fluctuates much more rapidly in time such that our assumption of a constant mean transcription initiation rate breaks down. We chose not to address this regime in this work because only a small minority of cells (4%) studied exhibited bursting behavior. Nevertheless, although our model does not capture bursting behavior (Section S4.3; Fig. S4E and F), transcriptional bursting remains a prevalent phenomenon in eukaryotic transcription and thus motivates extensions to this work to account for its behavior. For example, one possible implementation to account for transcriptional bursting could first utilize the widespread two-state model used to describe this phenomenon (***Peccoud and Ycart, 1995***) in order to partition a time trace into ON and OFF time windows. Then the MCMC inference method developed in this work could be used to quantify the transcription cycle during the ON and OFF windows with finer precision.

### Outlook

To conclude, while we demonstrated this inference approach in the context of the regulation of a *hunchback* reporter in *Drosophila melanogaster*, it can be readily applied to other genes and organisms in which MS2 and PP7 have been already implemented (***Golding et al., 2005***; ***Chubb et al., 2006***; ***Darzacq et al., 2007***; ***Garcia et al., 2013***; ***Lucas et al., 2013***; ***Tantale et al., 2016***; ***Lee et al., 2019***; ***Sato et al., 2020***), or where non-genetically encoded RNA aptamer technologies such as Spinach (***Paige et al., 2011***; ***Sato et al., 2020***) are available. Thus, we envision that our analysis strategy will be of broad applicability to the quantitative and molecular *in vivo* dissection of the transcription cycle and its regulation across many distinct model systems.

## Methods and Materials

### DNA constructs

The fly strain used to express constitutive MCP-mCherry and PCP-eGFP consisted of two transgenic constructs. The first construct, MCP-NoNLS-mCherry, was created by replacing the eGFP in MCP-NoNLS-eGFP (***Garcia et al., 2013***) with mCherry. The second construct, PCP-NoNLS-eGFP, was created by replacing MCP in the aforementioned MCP-NoNLS-eGFP with PCP, sourced from ***Larson et al***. (***2011a***). Both constructs were driven with the *nanos* promoter to deliver protein maternally into the embryo. The constructs lacked nuclear localization sequences because the presence these sequences created spurious fluorescence puncta in the nucleus that decreased the overall signal quality (data not shown). Both constructs were incorporated into fly lines using P-element transgenesis, and a single stable fly line was created by combining all three transgenes.

The reporter construct P2P-MS2-lacZ-PP7 was cloned using services from GenScript. It was incorporated into the fly genome using PhiC31-mediated Recombinase Mediated Cassette Exchange (RMCE) (***Bateman et al., 2006***), at the 38F1 landing site.

Full details of construct and sequence information can be found in a public Benchling folder.

### Fly strains

Transcription of the *hunchback* reporter was measured by imaging embryos resulting from crossing *yw;MCP-NoNLS-mCherry,Histone-iRFP;MCP-NoNLS-mCherry,PCP-NoNLS-GFP* female virgins with *yw;P2P-MS2-LacZ-PP7* males. The *Histone-iRFP* transgene was provided as a courtesy from Kenneth Irvine and Yuanwang Pan.

### Sample preparation and data collection

Sample preparation followed procedures described in ***Bothma et al***. (***2014***), ***Garcia and Gregor*** (***2018***), and ***Lammers et al***. (***2020***). To summarize, embryos were collected, dechorinated with bleach and mounted between a semipermeable membrane (Lumox film, Starstedt, Germany) and a coverslip while embedded in Halocarbon 27 oil (Sigma). Excess oil was removed with absorbent paper from the sides to flatten the embryos slightly. Data collection was performed using a Leica SP8 scanning confocal microscope (Leica Microsystems, Biberach, Germany). The MCP-mCherry, PCP-eGFP, and Histone-iRFP were excited with laser wavelengths of 488 nm, 587 nm, and 670 nm, respectively, using a White Light Laser. Average laser powers on the specimen (measured at the output of a 10x objective) were 35 *μ*W and 20 *μ*W for the eGFP and mCherry excitation lasers, respectively. Three Hybrid Detectors (HyD) were used to acquire the fluorescent signal, with spectral windows of 496-546 nm, 600-660 nm, and 700-800 nm for the eGFP, mCherry, and iRFP signals, respectively. The confocal stack consisted of 15 equidistant slices with an overall z-height of 7 *μ*m and an inter-slice distance of 0.5 *μ*m. The images were acquired at a time resolution of 15 s, using an image resolution of 512 x 128 pixels, a pixel size of 202 nm, and a pixel dwell time of 1.2 *μ*s. The signal from each frame was accumulated over 3 repetitions. Data were taken for 355 cells over a total of 7 embryos, and each embryo was imaged over the first 25 min of nuclear cycle 14.

### Image analysis

Images were analyzed using custom-written software following the protocols in ***Garcia et al***. (***2013***) and ***Lammers et al***. (***2020***). This software contains MATLAB code automating the analysis of all microscope images obtained in this work, and can be found on a public GitHub repository. Briefly, this procedure involved segmenting individual nuclei using the Histone-iRFP signal as a nuclear mask, segmenting each transcription spot based on its fluorescence, and calculating the intensity of each MCP-mCherry and PCP-eGFP transcription spot inside a nucleus as a function of time. The Trainable Weka Segmentation plugin for FIJI (***Arganda-Carreras et al., 2017***), which uses the FastRandomForest algorithm, was used to identify and segment the transcription spots. The final intensity of each spot over time was obtained by integrating pixel intensity values in a small window around the spot and subtracting the background fluorescence measured outside of the active transcriptional locus. When no activity was detected, a value of NaN was assigned.

### Data Analysis

Inference was done using *MCMCstat*, an adaptive MCMC algorithm (***Haario et al., 2001, 2006***). Figures were generated using the open-source gramm package for MATLAB, developed by Pierre Morel (***Morel, 2018***). Generalized linear regression used in Fig. 5 utilized a normally distributed error model and was performed using MATLAB’s *glmfit* function. All scripts relating to the MCMC inference method developed in this work are available at the associated GitHub repository.

## Acknowledgements

We thank Sandeep Choubey, Antoine Coulon, Jane Kondev, Anders Sejr Hansen, Mustafa Mir, Rob Phillips, Manuel Razo-Mejia, and Matthew Ronshaugen for thoughtful comments on the manuscript. We also are grateful to Florian Jug, Nick Lammers, and Armando Reimer for their crucial work in developing the image analysis code used here.

This work was supported by the Burroughs Wellcome Fund Career Award at the Scientific Interface, the Sloan Research Foundation, the Human Frontiers Science Program, the Searle Scholars Program, the Shurl and Kay Curci Foundation, the Hellman Foundation, the NIH Director’s New Innovator Award (DP2 OD024541-01), and an NSF CAREER Award (1652236) (HGG), an NSF GRFP (DGE 1752814) (EE, MT), a UC Berkeley Chancellor’s Fellowship (EE), a KFAS scholarship (YJK), and an DoD NDSEG graduate fellowship (JL).

## Supplementary Information

### S1 Full Model

To predict MS2 and PP7 fluorescence traces, we utilized a simple model of transcription initiation, elongation, and cleavage. The entire model has the following free parameters:

- ⟨*R*⟩, the mean transcription initiation rate
- δ*R*(*t*), the time-dependent fluctuations in the transcription initiation rate around the mean ⟨*R*⟩
- *v*_*elon*_, the RNAP elongation rate
- *τ*_*cleave*_, the mRNA cleavage time
- *t*_*on*_, the time of transcription onset after the previous mitosis, where *t* = 0 corresponds to the start of anaphase
- MS2_*basal*_, the basal level of MCP-mCherry fluorescence
- PP7_*basal*_, the basal level of PCP-eGFP fluorescence
- *α*, the scaling factor between MCP-mCherry and PCP-eGFP arbitrary fluorescence units

Note that the fluctuations δ*R*(*t*) are independent for each time point, and exist to allow for a slight time dependence in the overall initiation rate. Thus, δ*R*(*t*) parameterizes a set of independent constant offsets in the overall loading rate at each time point.

First, the parameters ⟨*R*⟩, δ*R*(*t*), *t*_*on*_, *v*_*elon*_, and *τ*_*cleave*_ were used to generate a map *x*_*i*_(*t*) of the position of each actively transcribing RNAP molecule *i* along the body of the reporter gene, as a function of time. Although the model is represented with continuous time, the subsequent computational simulation used for the statistical inference relies on discrete timesteps. Thus, given a computational time step *dt, R*(*t*)*dt* RNAP molecules are loaded at time point *t* at the promoter *x* = 0, where

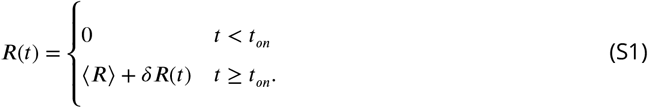

Note while *R*(*t*)*dt* is a floating point number, the model utilizes discrete numbers of RNAP molecules. As a result, *R*(*t*)*dt* is rounded down to the nearest integer since the. After initiation, each RNAP molecule proceeds forward with the constant elongation rate *v*_*elon*_. Once an RNAP molecule reaches the end of the gene, an additional cleavage time *τ*_*cleave*_ elapses after which the nascent transcript is cleaved and disappears instantly. This assumption of instantaneous disappearance following cleavage is justified in Section S3 based on the diffusion time scale of individual mRNA molecules. From this position map, and based on the locations of the stem loop sequences along the reporter construct (Fig. S1), we calculate the predicted MS2 and PP7 fluorescence signals. The contribution to the MS2 signal 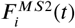 of an individual RNAP molecule *i* at position *x*_*i*_(*t*) is given by

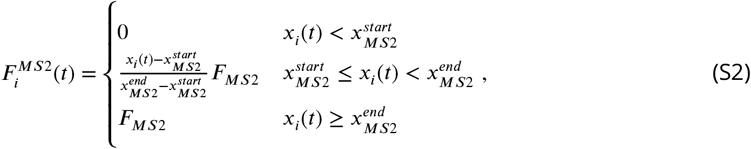

where 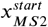 and 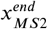 are the start and end positions of the MS2 stem loop sequence, respectively, and *F*_*MS*2_ is the mCherry fluorescence produced by a single RNAP molecule that has transcribed the entire set of MS2 stem loops. Here, we also assume that RNAP molecules that have only partially transcribed the MS2 stem loops result in a fractional fluorescence given by the fractional length of the MS2 stem loop sequence transcribed. Similarly, the contribution to the PP7 signal 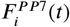 is given by

**Figure S1.**
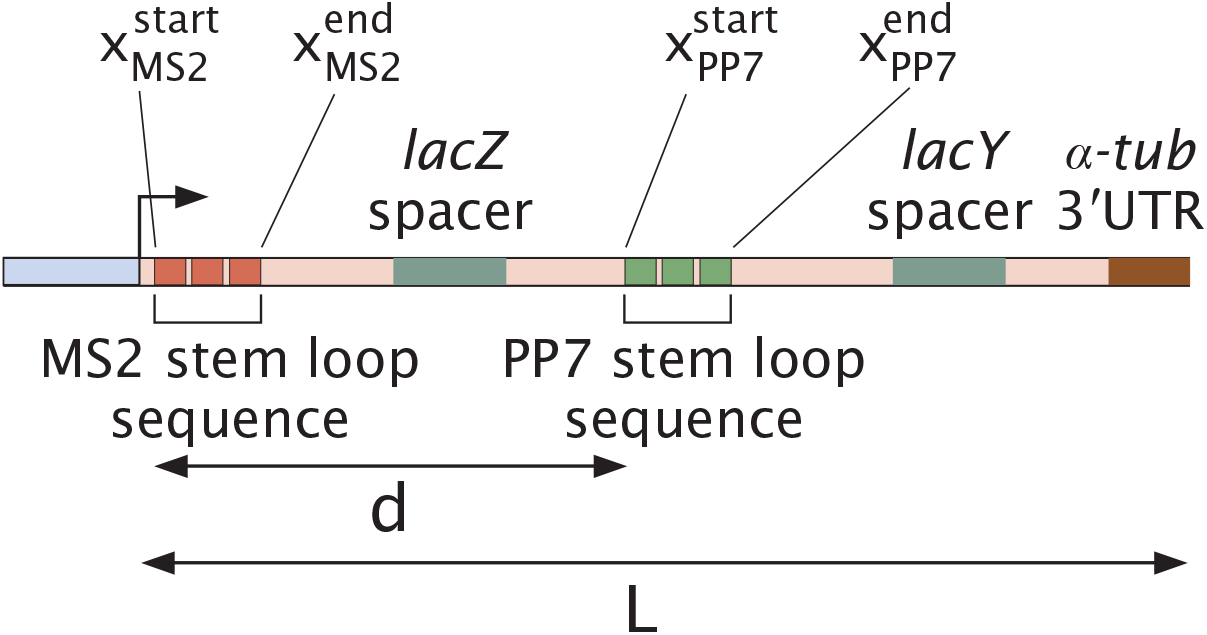
Detailed description of reporter construct used in this work. Labeled positions are 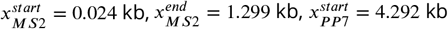, and 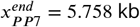, where *x* = 0 corresponds to the 3’ end of the promoter. Distances are *d* = 4.27 kb and *L* = 6.63 kb.

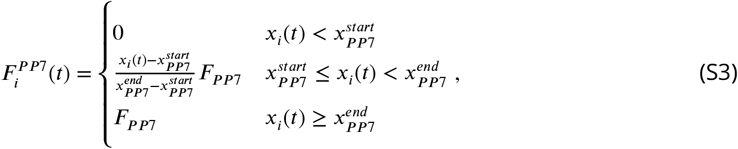

where 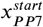 and 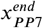 are the start and end positions of the PP7 stem loop sequence, respectively, and *F*_*PP*7_ is the GFP fluorescence produced by a single RNAP molecule that has transcribed the entire set of PP7 stem loops. Note that we assume that the MCP-mCherry and PCP-GFP fluorophores effectively bind instantaneously to all their associated stem loops once they are transcribed. Due to the high numbers of nascent transcripts on the reporter gene (Fig. 5D), we expect that corrections to this assumption due to incomplete, stochastic, and/or non-instantaneous fluorophore binding will not introduce substantial deviations to the model.

The temporal dynamics of the total MS2 and PP7 signals *F*_*MS*2_(*t*) and *F*_*PP*7_(*t*) are then obtained by summing over all the individual RNAP molecule contributions for each timepoint

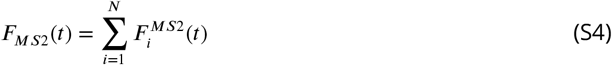

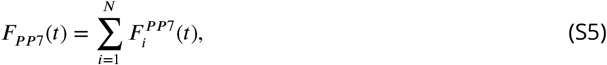

where *i* is the index of each individual RNAP molecule and *N* is the total number of loaded RNAP molecules. The final signal is then modified by accounting for the scaling factor e*α* and the basal fluorescence values of MS2_*basal*_ and PP7_*basal*_. e*α* is necessary because the two fluorescent protein signals have different arbitrary units (Fig. 3). Further, the two basal fluorescence values are incorporated to account for the experimentally observed low baseline fluorescence in each fluorescent channel. The final signals 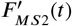 and 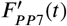 are then given by

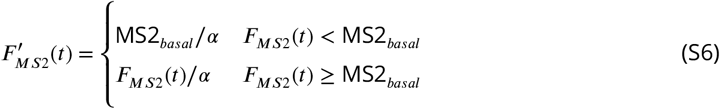

and

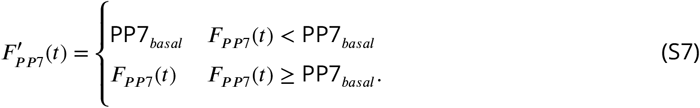

All of the model parameters introduced in this section were used as free parameters in the fitting procedure described in Section S4.

Note that the model does not make mechanistic claims about the nature of the cleavage process, which could potentially be convolved with processes such as transcriptional pausing. Specifically, if RNAP pausing were to happen 3’ of the PP7 stem loop sequence, then it is effectively indistinguishable from cleavage at the 3’ UTR.

However, we stress that our model is only an effective parameterization, and so we make no mechanistic claims as to the source of a particular cleavage time value. What our model interprets as cleavage could stem from pausing at the 3’UTR of the reporter, for example, or from continued elongation past the 3’UTR due to inefficient cleavage and termination processes. These would exhibit the same experimental signals—namely, persistence of fluorescent signal after the expected time of signal loss—and thus is a challenge of experimental resolution and not of model formulation.

## S2 Characterization of photobleaching in experimental setup

To determine whether photobleaching was present in our experimental setup, we conducted an experiment with the dual-color 5’/3’ tagged reporter (Fig. 1C) where half of the field of view was illuminated using the experimental settings described in the Materials and Methods section (Fig. S2A, purple), and the other half was illuminated at half the temporal sampling rate (Fig. S2A, yellow).

Since the measurement conditions were identical except for the sampling rate for both reporter constructs used in this work, any systematic differences between the two measurement conditions could only stem from this different sampling rate. Thus, if the experimental settings were in the photobleaching regime, then the purple region would exhibit fluorescence at a systematically lower intensity compared to the yellow region. Figures S2B and C shows the fluorescence intensities of mCherry and eGFP as a function of time at a particular anterior-posterior position of the embryo for both 0.5x and 1x sampling rates, where data points indicate fluorescence averaged within the anterior-posterior position (indicated schematically by the dashed box in Fig. S2A) and error bars indicate standard error across cells. The plots reveal that, qualitatively, there is no obvious systematic difference between the two illumination regions.

To quantify photobleaching, we defined the average normalized difference Δ between illuminated regions. This magnitude is calculated by subtracting the fluorescence value at 1x sampling rate *F*_1*x*_ by that at 0.5x sampling rate, dividing by the fluorescence value at 0.5x sampling rate *F*_0.5*x*_, and then averaging across all time points *N*_*timepoints*_ and embryo positions *N*_*positions*_

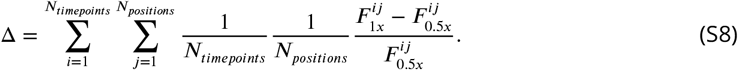

For example, for the curves shown in Fig. S2B, this entails subtracting the red curve by the black curve, dividing by the black curve, and then averaging for all anterior-posterior embryo positions. An overall value of less than zero means that the 1x sampling rate produces systematically lower fluorescence intensities, indicating that our experimental settings are in the photobleaching regime. As seen in Figure S2D, the average normalized difference Δ is consistent with zero for both fluorophores (within standard error, measured across all time points and anterior-posterior positions). Thus, we conclude that our data are not in the photobleaching regime.

**Figure S2.**
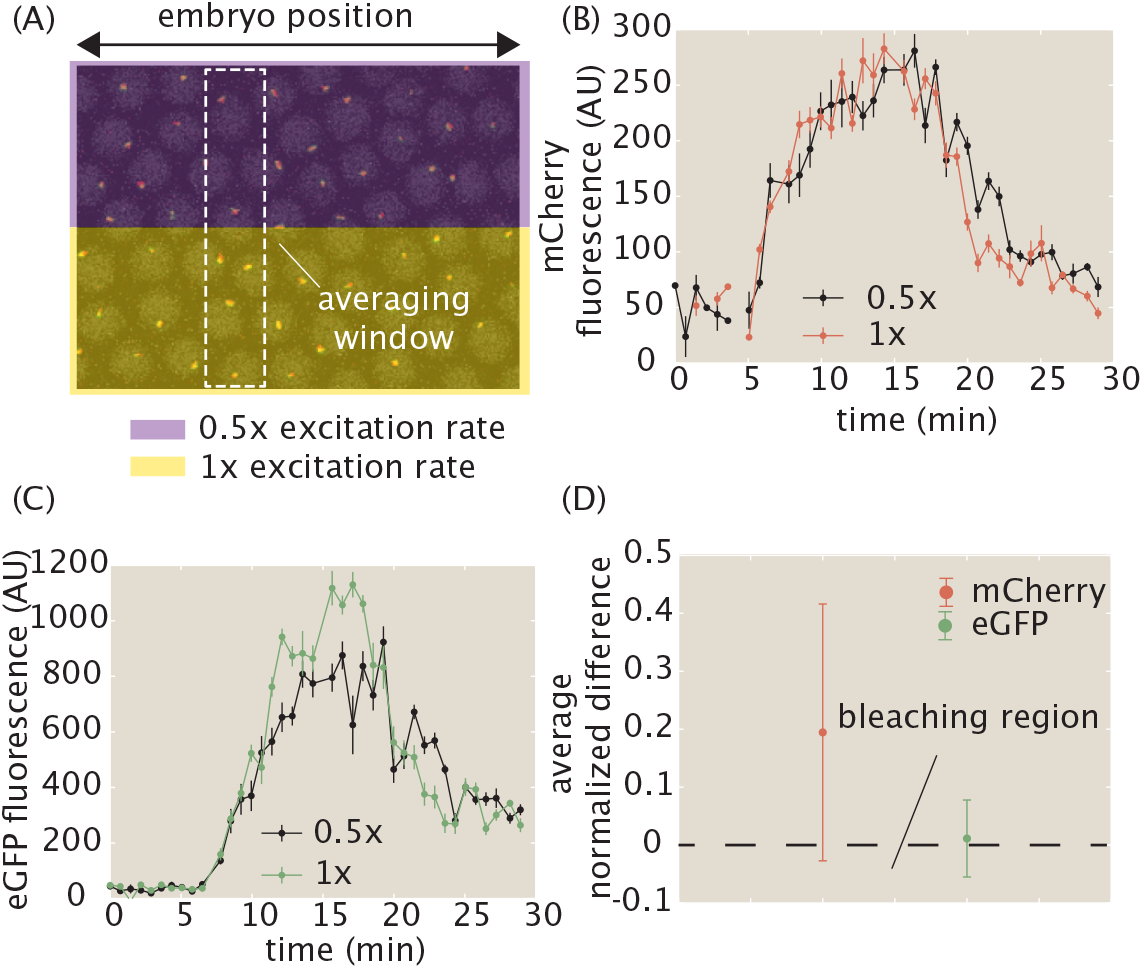
Investigation of photobleaching in experimental setup. (A) Control experiment where half of the field of view is illuminated at the standard experimental settings (yellow), and the other half of the field of view is imaged at half of the illumination rate (purple). (B, C) The (B) mCherry and (C) eGFP fluorescence signals at a given anterior-posterior embryo position, averaged across cells within that position (white dashed rectangle in (A)), do not exhibit photobleaching. (D) The average normalized difference between illuminated regions, averaged across time points and anterior-posterior embryo positions, are approximately zero within error. A negative value would indicate the presence of photobleaching. (B, C, error bars indicate standard error of the mean averaged across cell nuclei in the field of view; D, error bars indicate standard error of the mean averaged across time points and embryo positions).

## S3 Justification for approximating transcript cleavage as instantaneous

In the model presented in Section S1, we assumed that, when a nascent RNA transcript is cleaved at the end of the reporter gene, its MS2 and PP7 fluorescence signals disappear instantaneously. Here, we justify this assumption by demonstrating that the timescale of mRNA diffusion away from the active locus is much shorter than the experimental resolution of our system.

When a nascent RNA transcript is cleaved, it diffuses away from the gene locus. For a free particle with diffusion coefficient *D*, the characteristic timescale *τ* to diffuse a length scale *L* is given by

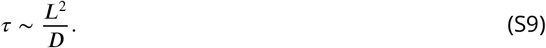

In the context of the experiment performed here, this can interpreted as the timescale for a cleaved mRNA transcript to diffuse away from the diffraction-limited fluorescence punctum at the locus.

We can estimate the characteristic timescale *τ* by plugging in the following values. Assume that the completed transcript possesses a typical mRNA diffusion coefficient of *D* ∼ 0.1 *μm*^2^/*s* (***Gorski et al., 2006***). The length scale *L* corresponds to the Abbe diffraction limit, which yields *L* ∼ 250 *nm* for green light with a wavelength of about 500 *nm* and a microscope with a numerical aperture of 1. Plugging these values into the equation yields a diffusion time scale of

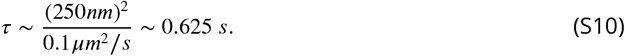

As a result, a newly cleaved mRNA transcript will typically diffuse away from the locus in less than a second, meaning that its MS2 and PP7 fluorescence signal will vanish much faster than our experimental time resolution of 15 *s*. For this reason, we can justify approximating the cleavage process as instantaneously removing the fluorescent signals of newly cleaved transcripts.

## S4 MCMC inference procedure

### S4.1 Overview and application of MCMC

The inference procedures described in the main text were carried out using the established technique of Markov Chain Monte Carlo (MCMC). Specifically, we used the MATLAB package MCMCstat, an adaptive MCMC technique (***Haario et al., 2001, 2006***). For detailed descriptions, we refer the reader to the the MCMCstat website (https://mjlaine.github.io/mcmcstat/), as well as to a technical overview of MCMC (***Geyer, 1992***). Briefly, MCMC allows for an estimation of the parameter values of a model that best fit the experimentally observed data along with an associated error. In this work, we use MCMC to infer the best fit values of the transcription cycle parameters given observed fluorescence data at the single-cell level. Then, we combine these inference results across cells to construct distributions of inferred values across the ensemble of cells.

MCMC calculates a Bayesian posterior probability distribution of each free parameter given the data by stochastically sampling different parameter values. For a given set of observations *D* and a model with parameters *θ*, the so-called posterior probability distribution of *θ* possessing a particular set of values is given by Bayes’ theorem

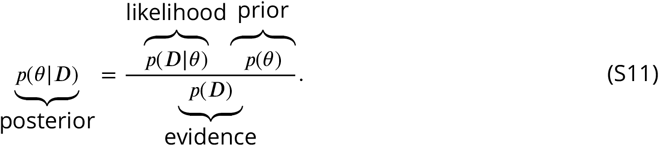

This posterior distribution is a combination of three components: the likelihood, prior, and evidence. This latter term represents the probability of the observations possessing their particular values, and allows the overall posterior distribution to be normalized. In practice, the evidence term is often dropped since MCMC can still yield accurate results without requiring this normalization. Thus, we have

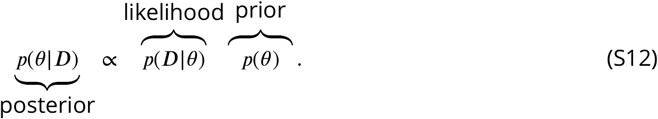

The prior function contains *a priori* assumptions about the probability distribution of parameter values *θ*, and the likelihood function represents the probability of obtaining the observations, given a particular set of parameters *θ*. Thus, the *most likely* set of parameters *θ* occurs when the product of the likelihood and prior is maximized, resulting in a maximum in the posterior function. MCMC extends this by sampling different values of *θ* such that an approximation of the full posterior distribution is also obtained.

The prior distributions for the inferred parameters were set as follows. The prior distribution for the fluctuations in the initiation rate δ*R*(*t*) at each time point was assumed to be a Gaussian distribution centered around 0 AU/min with a standard deviation of 30 AU/min. This penalized fluctuations that strayed too far from zero, smoothing the overall initiation rate *R*(*t*). For the rest of the parameters, a uniform distribution was chosen using the following uniform intervals:

- v_*elon*_:[0, 10] kb/min
- *t*_*on*_:[0, 10] min
- *α*:[0, 1]
- *τ*: [0, 20] min
- MS2_*basal*_:[0, 50] AU
- PP7_*basal*_:[0, 50] AU
- 〈*R*〉:[0, 40] AU/min

These intervals were justified with the following arguments. Previous elongation rate measurements have indicated values between around 1 and 4 kb/min (Fig. S9; (***Ardehali and Lis, 2009***)), so we approximately doubled this range for flexibility. Previous measurements of the transcription onset time *t*_*on*_ for *hunchback* range from about 1 to 6 min (***Garcia et al., 2013***), so we chose a similarly flexible interval. The calibration factor e*α* must take on values between 0 and 1, since, under the experimental settings used, mCherry exhibits weaker absolute fluorescence than eGFP (see for example, Fig. 3C). Although the cleavage time is not well understood, estimates lie on the order of minutes (***Lenstra et al., 2016***)—we chose a large interval to be conservative. Based on our experimental data (e.g. Fig. 2B), basal levels of MS2 and PP7 fluorescence lie comfortably in the range [0, 50] AU. Finally, as observed in our data and also reported in ***Garcia et al***. (***2013***), the mean rates of initiation lie comfortably in the range [0, 40] AU/min (Fig. 4A).

For the likelihood function, a Gaussian error function was used

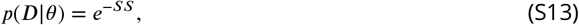

where SS is a scaled sum-of-squares residual function given by

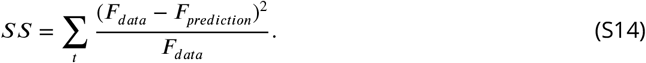

Here, the summation runs over individual time points, *F*_*data*_ corresponds to the MS2 or PP7 fluorescence at a given timepoint, and *F*_*prediction*_ corresponds to the predicted MS2 or PP7 fluorescence according to the model, for a given set of parameter values. That is,

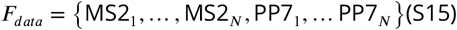

where the subscripts indicate the time index over *N* time points. Similarly,

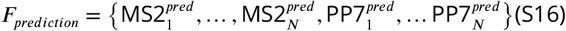

where the superscripts indicate that these are model predictions evaluated at the experimental time points. The presence of *F*_*data*_ in the denominator scales the overall sum-of-squares residual function by the mean signal intensity and is required because the measurement noise in the fluorescence scales linearly with fluorescence intensity (Section S4.2 and Fig. S3).

The MCMC approach samples values of parameters *θ* to approximate the posterior probability distribution. There are several algorithms that achieve this—the adaptive technique used in the MCMCstat package is an efficient algorithm that updates the sampling technique to more quickly arrive at the converged distribution.

For each inference run, an initial condition of parameter values is chosen. The algorithm then stochastically updates the next set of parameter values based on the current and previous values of the posterior distribution function. After a preset number of updates (typically at least on the order of thousands), the algorithm stops, resulting in a *chain* of MCMC parameter value samples. The initial period following the initial condition, known as the *burn-in* time, is typically discarded since the results are not reliable. The remaining values of the chain comprise an approximation of the underlying posterior probability distribution, with smaller errors for longer run times.

For the purposes of this work, the MCMC procedure was run by separately inferring parameter values for the data corresponding to each single cell. For each inference, random parameter values were chosen for the initial condition of the sampling algorithm in order to prevent initial condition bias from affecting the inference results. The algorithm was run for a total of 20, 000 iterations, which, after removing a burn-in window of length 10, 000, resulted in a chain of length 10, 000 for each of the 355 cells examined. To assess whether or not the algorithm was run for a sufficient number of iterations, the final chain was examined for *rapid mixing*, where the sampled values of a particular parameter rapidly fluctuate around a converged value. Figure 2C highlights this rapid mixing in the inferred transcription cycle parameters of a sample single cell. The lack of long-timescale correlations, also exemplified by the quick decay of the auto-correlation function of each chain (Fig. 2D), indicates that the algorithm has converged. In addition, a corner plot of the three transcription cycle parameters (Fig. 2E) illustrates the pairwise correlations between them, demonstrating that the inference did not encounter degenerate solutions, and that each parameter has a fairly unimodal distribution.

These diagnostics provided a check on the quality of the inference results. Afterwards, the mean value of each parameter’s final chain was then retained for each single cell for use in the further statistical analysis carried out in the main text.

### S4.2 Justification of scaled observation model due to fluorescence noise behaviour

The observation model parameterized by the sum-of-squares residual in Equation S14 is scaled by dividing by the overall fluorescence intensity. This is needed because the fluorescence noise is not constant, but rather scales linearly with overall intensity. Here, we demonstrate this behavior by examining the fluorescence noise exhibited in our system.

*A priori*, if we consider that the fluorescent signals in our experiment are the result of the sum of many individual fluorophores, then we would expect that, if an individual fluorophore possesses some intrinsic constant measurement error with variance *σ*^2^, then the associated error of *N* fluorophores would have a similarly scaled overall measurement error with variance *Nσ*^2^. Since *N* is proportional to the overall mean fluorescent signal, the observation model in Equation S14 thus needs the mean signal in the denominator.

To validate this scaling of the variance with the mean, we examined the data from the dual-color interlaced MS2/PP7 reporter construct from Figure 3B. These data constitute, in principle, a two-point measurement of the same underlying biological process, so we reasoned that we could utilize this measurement to quantify the scaling of fluorescence noise with respect to overall fluorescence intensity.

**Figure S3.**
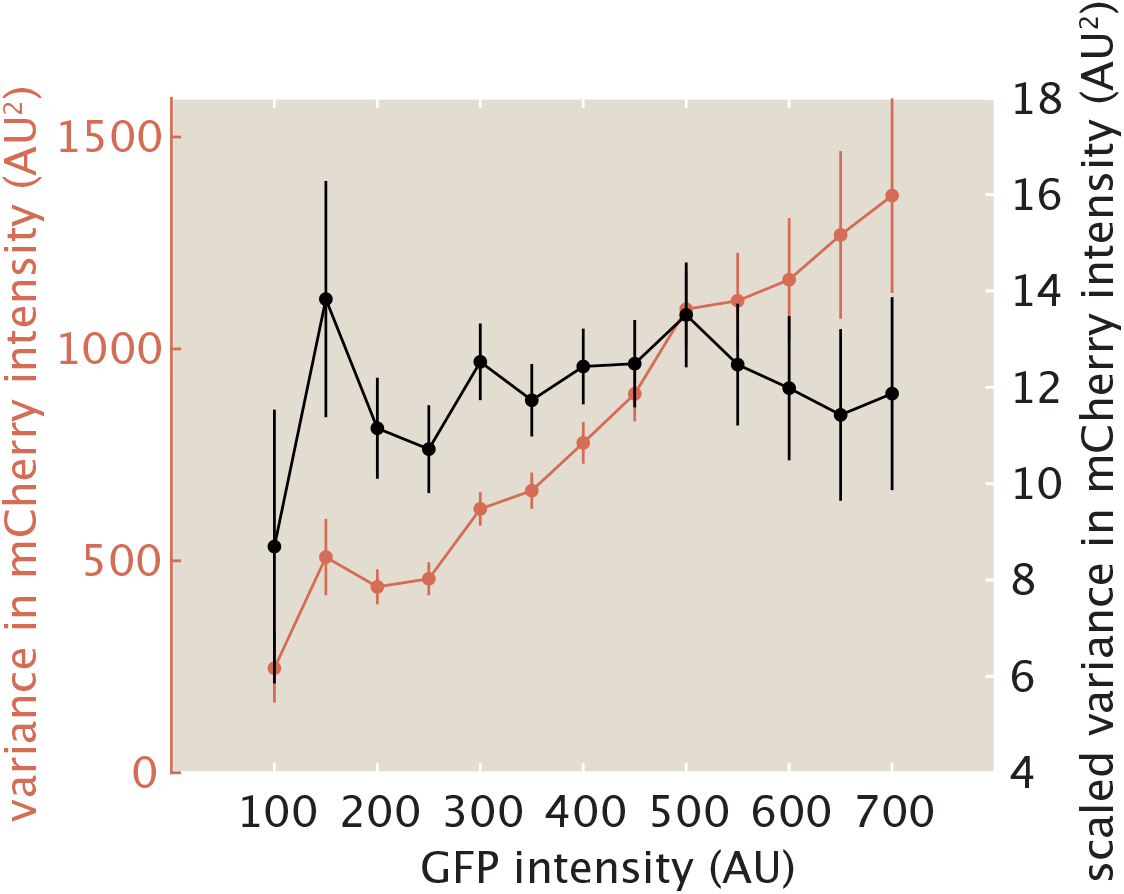
Scaling of fluorescence measurement noise with overall fluorescence intensity. Variance of mCherry fluorescence at a particular GFP fluorescence (red), from the dual-color interlaced reporter construct from Figure 3B, along with variance scaled by dividing out the mean mCherry fluorescence (black).

Specifically, by creating bins of eGFP fluorescence measurement from the scatterplot in Figure 3D, we calculated how the variance of associated mCherry fluorescence values within a bin scaled with eGFP fluorescence (here a proxy for overall fluorescence intensity). If the calculated variance increased with overall fluorescence, this would indicate that the fluorescence measurement noise is not constant, but rather scaled positively with signal strength. Figure S3 shows this calculated variance (red), along with bootstrapped standard error, as a function of bin value (i.e. eGFP fluorescence). We see that the variance indeed increases with bin value fairly linearly, confirming our hypothesis. If we then scale the variances by dividing by the mean mCherry fluorescence within a bin, we recover a constant scaling, as expected (black).

### S4.3 Curation of inference results

Individual single cell inference results were filtered automatically and then run through an automated curation procedure for final quality control. First, due to experimental and computational imaging limits, some MS2 or PP7 trajectories were too short to run a meaningful inference on. As a result, we automatically skipped over any cell with an MS2 or PP7 signal with fewer than 30 datapoints. This amounted to 626 cells skipped out of a total of 1053, with 427 (41%) retained.

Second, the retained cells were run through an automated curation pipeline. For each single-cell fit, we calculated the average squared normalized residual δ^2^, defined as

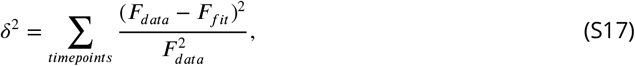

where the summation occurs over all time points and *F*_*data*_ and *F*_*fit*_ correspond to the fluorescence data and fit, respectively. Thus, δ^2^ gives a measure of how good or bad, on average, each single-cell fit is. Figure S4A and B show histograms of the average squared normalized residual δ^2^ for the entire *n* = 427 dataset, with log and linear x-axes. We see that the vast majority of data possesses values of δ^2^ smaller than unity, with a long tail at higher values corresponding to bad fits. We decided to implement a cutoff of 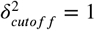 (red line), where any cell with a higher value of δ^2^ was automatically discarded.

In sum, 355 cells of data were retained out of 427 total after this curation process. We reasoned that, since we still ended up with hundreds of single cells of data, the resultant statistical sample size was large enough to extract meaningful conclusions.

To assess the rejected fits for underlying biological causes, we did a qualitative examination for common features. There were several sources of bad fits. First, some traces possessed low signal-to-noise ratio (Fig. S4C) that nevertheless yielded reasonable fits that were slightly above 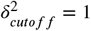. Still others simply had poor fits, possibly due to running into issues with the inference algorithm such as getting trapped in local minima (Fig. S4D). We consider improvements to the algorithm to be outside the scope of this work, since the retained data still contain enough statistical size to provide interpretable results.

Finally, one potential biological source confounding the model could be the presence of substantial transcriptional bursting of the promoter. Although the majority of the traces we analyzed indicated that the *hunchback* reporter gene studied here possessed a promoter that was effectively ON during the cell cycle studied, a small fraction of traces (4% of the filtered cells) possessed substantial time dependence of the fluorescence signal, potentially resulting from rapid switching of the promoter between ON and OFF states (Fig. S4E).

The presence of transcriptional bursts is of high biological significance, but capturing the behavior would require more specific models (e.g. two-state telegraph models like ***Lammers et al***. (***2020***)). As a result, we relegate extensions of the model that can account for transcriptional bursting for future work. Thus, our work provides a self-contained framework applicable for describing the behavior of promoters that are primarily ON for the duration of the experiment and that do not experience transcriptional bursting.

Due to the variety of sources contributing to the rejected fits, we opted for a conservative approach and only analyzed the cells with high signal quality that did not exhibit the complications mentioned above. The number of retained fits were still much higher than the number of rejected fits (Fig. S4F).

To check that the curation procedure did not incur substantial bias, we compared the average inferred mean initiation rate, elongation rate, and cleavage time as a function of embryo position between the post-filtering curated and uncurated datasets of size *n* = 355 and *n* = 427, respectively (Fig. S4G-I). We observed no substantial difference between the two datasets, indicating that the curation procedure was not systematically altering the inference results.

### S4.4 Validation of inference results

To assess the accuracy of the inference method, we validated our MCMC approach against a simulated dataset. Using the inferred distribution of model parameters from the experimental data, we generated a simulated dataset with our theoretical model (Section S1) and ran the MCMC inference on it.

The simulated dataset consisted of 300 cells. The model parameters used to simulate each individual cell’s MS2 and PP7 fluorescences were drawn randomly from a Gaussian distribution, with mean *μ* and standard deviation *σ* calculated from the distribution of inferred model parameters from the experimental data. Table S1 shows the parameters used in the Gaussian distributions generating each single cell’s model parameters. We chose to fix the time-dependent fluctuations in the initiation rate δ*R*(*t*) at zero since these fluctuations are not well understood at the single-cell level, and the *hunchback* reporter studied here is well parameterized by a mean initiation rate (Fig. 2B).

**Figure S4.**
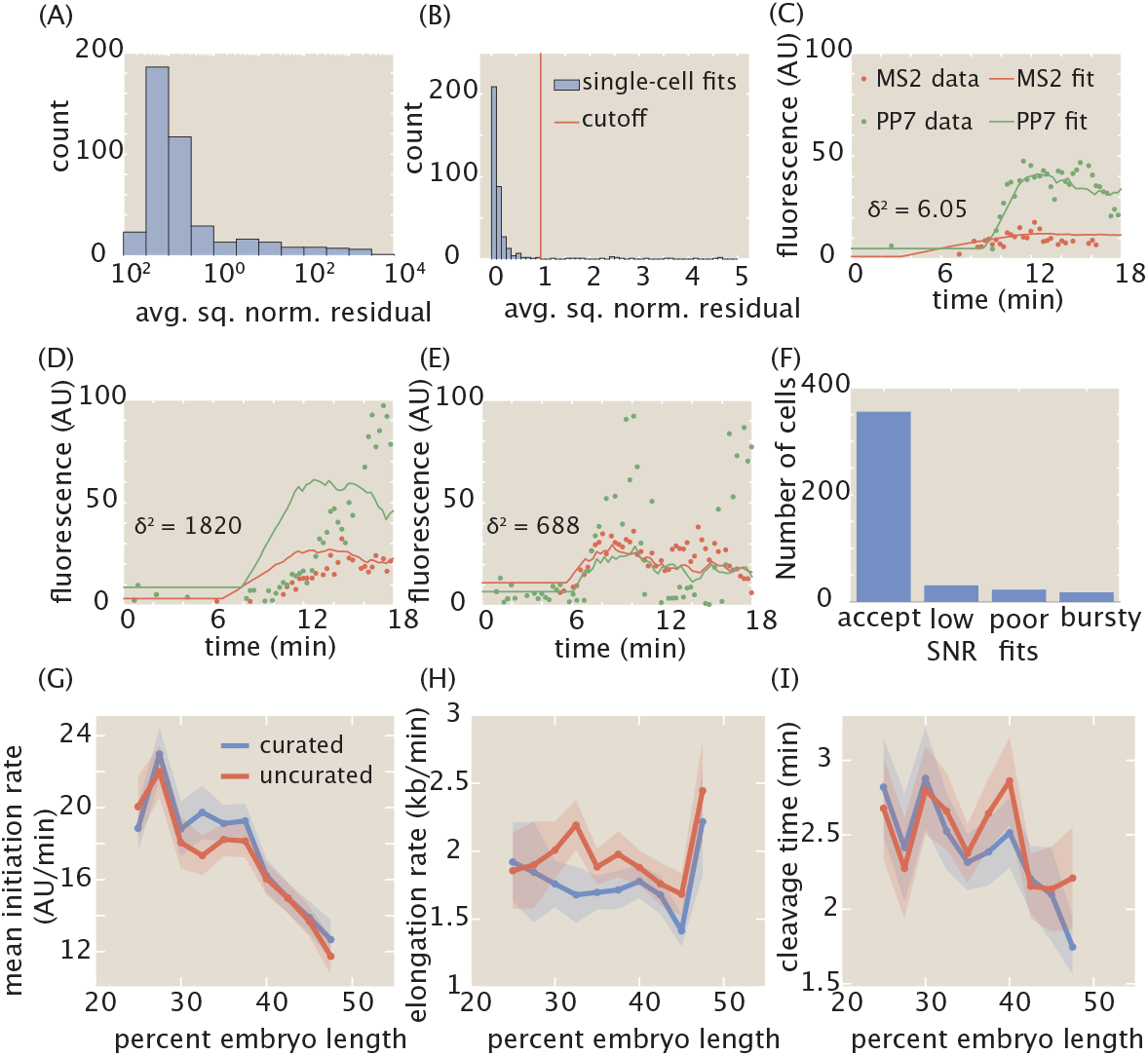
Automated curation of data. (A, B) Histograms (blue) of average squared normalized residual of single-cell fits, in log (A) and linear (B) scale, with cutoff of 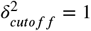 shown in red in (B). (C) Example of bad fit from poor signal-to-noise ratio (SNR). (D) Example of bad fit of otherwise reasonable data from issues in fitting algorithm, for example due to local minima. (E) Example of bad fit due to potential presence of substantial bursting of promoter. (F) Number of single cell fits in each class of rejected fit, along with number of accepted fits, after the initial filtering based on number of time points. Altogether, 84% of filtered fits were accepted. The percentages of filtered fits in the three rejected categories (low SNR, poor fits, bursting) were 7%, 5%, and 4%, respectively. The data shown in C-E are in each fluorophore’s intrinsic arbitrary unit without rescaling, to present the fluorescence intensities in their raw form. (G, H, I) Comparison of average inferred (G) mean initiation rate, (H) elongation rate, and (I) cleavage time as a function of embryo position, between curated (blue) and uncurated (red) datasets. Values of δ^2^ were 6.05, 1820, and 688 for the example fits shown in C-E, respectively, here given to illustrate the qualitative correspondence of δ^2^ as a metric with the overall goodness-of-fit. Shading in G-I represent standard error of the mean for 355 and 427 cells across 7 embryos for curated and uncurated datasets, respectively.

In addition, fluorescence measurement error was generated for each single cell and at each time point by drawing a random number from a Gaussian distribution with mean 0 and standard deviation 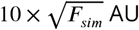 AU, where *F*_*sim*_ is the fluorescence at each time point, and adding this random number to the MS2 or PP7 fluorescence at that time point (prior to rescaling the MS2 fluorescence with the scaling factor e*α*). Here, the 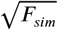 factor in the magnitude of the fluorescence noise accounts for our observation that the variance of the fluorescence measurement noise scales linearly with the mean signal intensity (Fig. S3).

Figure S5A shows an example of the simulated MS2 and PP7 fluorescence from a single cell along with their corresponding fits. The resulting MCMC-sampled values of the mean initiation rate, elongation rate, and cleavage time are shown in the histograms in Figure S5B (blue), along with the ground truth for that single cell (red line). As described in Section S4.1, the mean value of each sampled distribution was retained for downstream statistical analysis.

The accuracy of the inference was investigated on three levels: 1) systematic errors affecting mean analyses, 2) random errors affecting measurements of distributions, and 3) spurious correlations between parameters affecting inter-parameter correlations.

**Figure S5.**
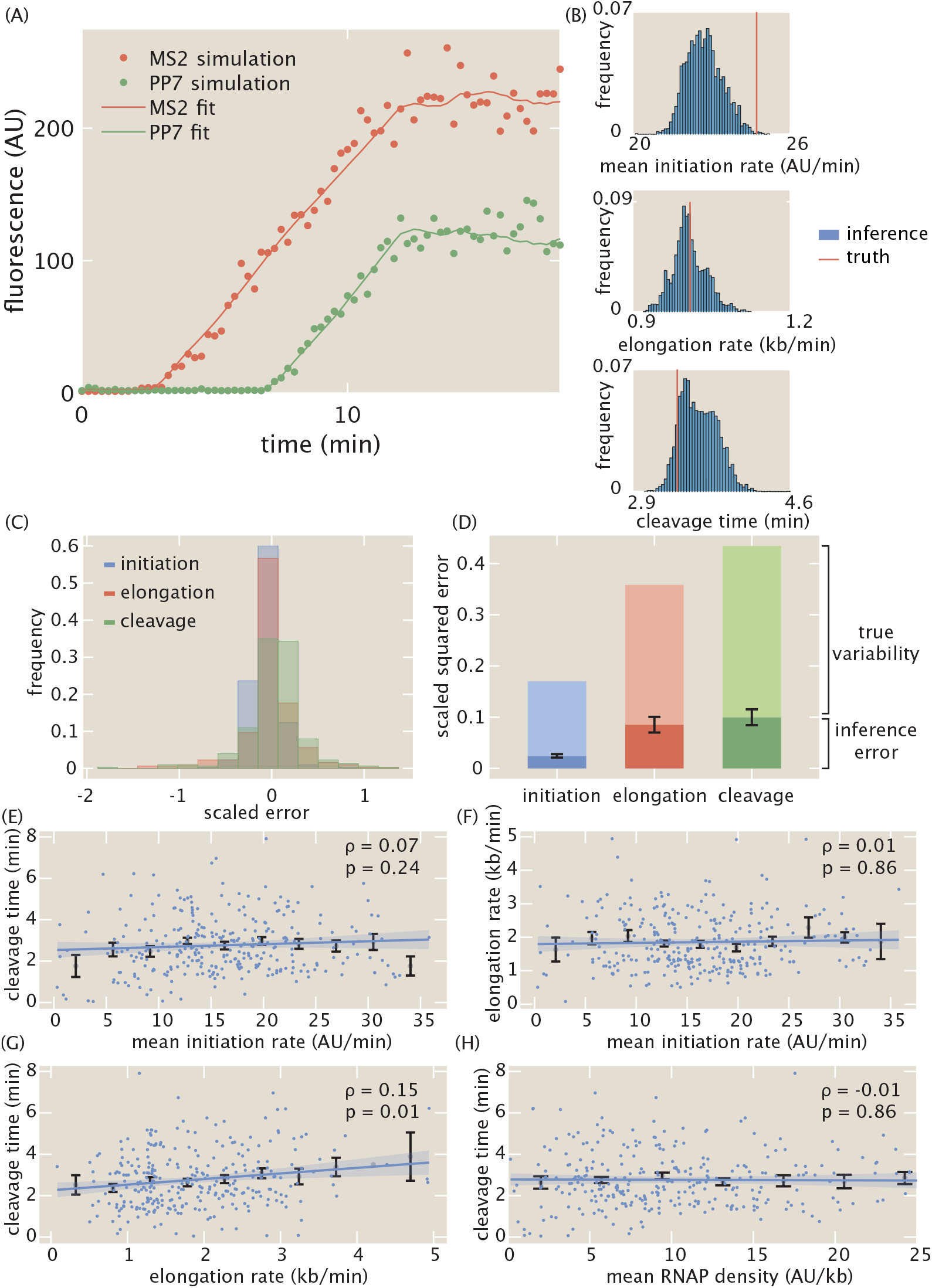
Overview of MCMC inference validation. (A) Example single-cell simulated data and inferred fits. (B) MCMC inference results for the simulated data in (A) for the mean initiation rate, elongation rate, and cleavage time. The histogram represents the raw MCMC sampled values, and the red line is the ground truth for this particular cell. The mean value of each histogram is then retained for further statistical analysis. (C) Scaled error of initiation, elongation, and cleavage for each simulated cell. (D) Comparison of relative magnitudes of random inference error and true experimental variability for the initiation, elongation, and cleavage parameters. (E, F, G, H) Single-cell correlations along with Spearman correlation coefficients and p-values for simulated data between (E) mean initiation rate and cleavage time, (F) mean initiation rate and elongation rate, (G) elongation rate and cleavage time, and (H) mean RNAP density and cleavage time, respectively. Blue points indicate single-cell values; black points and error bars indicate mean and SEM, respectively, binned across x-axis values. Line and shaded region indicate generalized linear model fit and 95% confidence interval, respectively. Linear fits were calculated using a generalized linear regression model and are presented for ease of visualization (see Materials and Methods for details).

**Table S1.**
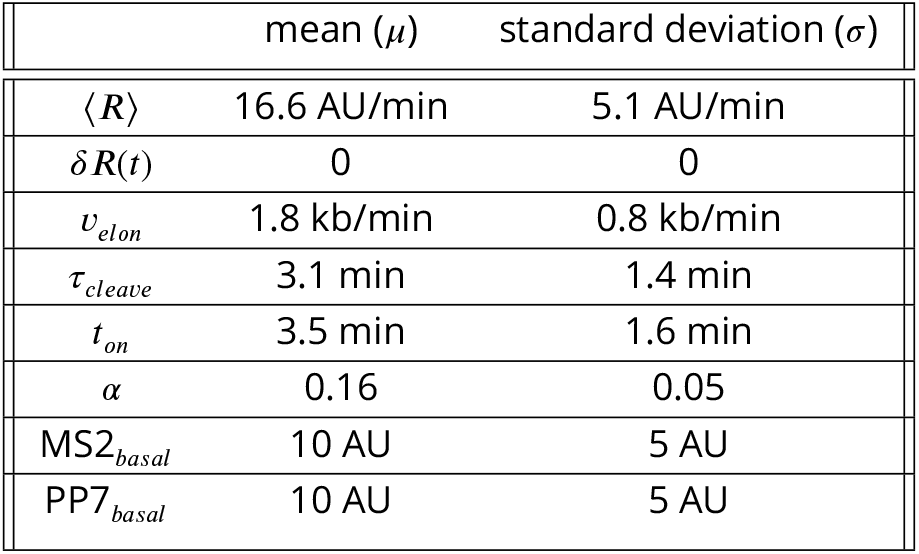
Mean and standard deviation of model parameters used in single-cell simulations.

| | mean ($\mu$) | standard deviation ($\sigma$) |
| --- | --- | --- |
| $\langle R \rangle$ | 16.6 AU/min | 5.1 AU/min |
| $\delta R(t)$ | 0 | 0 |
| $v_{elon}$ | 1.8 kb/min | 0.8 kb/min |
| $\tau_{cleave}$ | 3.1 min | 1.4 min |
| $t_{on}$ | 3.5 min | 1.6 min |
| $\alpha$ | 0.16 | 0.05 |
| MS2<sub>basal</sub> | 10 AU | 5 AU |
| PP7<sub>basal</sub> | 10 AU | 5 AU |

First, the scaled error *ε* for each parameter was calculated on a single-cell basis as defined by

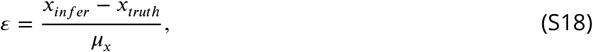

where *x* represents the model parameter being investigated, the subscripts indicate whether the quantity is the inferred result or the ground truth for that single cell, and *μ*_*x*_ is the population mean of the parameter value from the experimental data (i.e., the values of the “mean” column in Table S1). For example, for the mean initiation rate *R, μ*_⟨*R*⟩_ takes the value 16.6 AU/min. *ε* gives a unitless measure of the magnitude of inference error of each single cell, where a value of 1 indicates an error that is as large as the population mean itself. Because the scaled error is defined as the error due to inference for a single cell, it is an intensive quantity that is independent from the overall dataset size.

Figure S5C shows the histogram of single-cell scaled errors 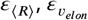, and 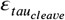 for the inferred mean initiation rate, elongation rate, and cleavage time, respectively. The majority of the scaled errors fall between −0.5 and 0.5, indicating that most inferred results possess relatively small error.

The systematic error on measurements of the ensemble mean can be estimated by calculating the mean of the scaled errors shown in Figure S5C. Doing so results in a value of −0.06 ± 0.01, −0.01 ± 0.02, and 0.04 ± 0.02 (mean and SEM) for the mean scaled error of the mean initiation rate, elongation rate, and cleavage time, respectively. For context, this means that, if the mean cleavage time is ∼ 3 min, then the systematic error in the cleavage time is ∼ 10 sec, about the time resolution of the data. Thus, the systematic error for each parameter is a couple orders of magnitude below that of the experimental mean value of each parameter, indicating that the inference provides an accurate and precise readout of the mean.

While the inference’s systematic error across cells may be small, the presence of individual single-cell errors will affect measurements of distributions of parameters. To investigate the impact of these random errors, we quantified the fraction of total empirically inferred variability that consisted of inferential error. Specifically, for a parameter *x*, we separated the variance of single-cell measurements as

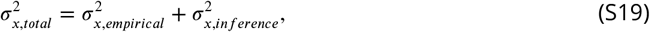

where 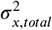 represents the overall single-cell variability observed in the data (the combination of empirical and inferential variability), 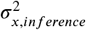 represents the error inherent to our inference process, and 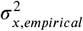 represents the true empirical variability after subtracting out inferential error 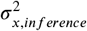. Note that 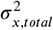 is the square of the values in the standard deviation column in Table S1.

Dividing by the square of the population means *μ*_*x*_ yields

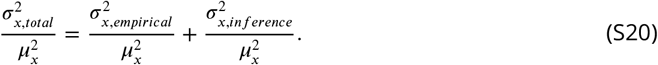

Note that these are just squared CV terms, and that the last term is simply the square of the scaled error *ε* defined earlier

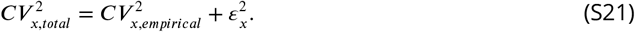

Thus, the overall impact of the inferential error can be quantified by calculating the relative magnitudes of the contributions of 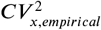 and *ε*^2^ to the total variability 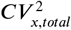. Figure S5D shows this separation, where the dark bars represent the squared scaled error *ε*^2^, the light bars represent the true empirical variability 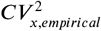, and the overall bars represent the total variability 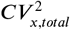 obtained from the values of *μ* and *σ* in Table S1.

All three model parameters—initiation, elongation, and cleavage—possess no more than approximately 25% inferential error. Nevertheless, the presence of this much error indicates that measurements of distributions of these parameters will be somewhat confounded by the inherent error present in our inference method, highlighting the general difficulty in measuring values beyond the mean.

However, these errors in the inference of the variability of the transcription cycle parameters should not impact the results of investigating the distribution of elongation rates in Figure 4D, since the simulated results there were also pushed through the inference pipeline and should pick up similar inferential noise. Furthermore, the variances of the simulated distributions in the presence or absence of single-molecule elongation variability differed by essentially around a factor of two (Fig. S10D), twice as much as the random error exhibited in the simulated results here (see Section S10 for details).

Future improvements on increasing the accuracy of measurements of distributions could be achieved, for example, by utilizing interleaved loops such as those introduced in Figure 3B. Here, two orthogonal species of mRNA binding proteins fused to different fluorescent proteins would bind to interleaved loops located at the 5’ end of the construct. In addition, a second pair of mRNA binding proteins would bind to an analogous set of interleaved loops located at the 3’ end. The result would be a four-color experiment, with two colors reporting on transcription at the 5’ end of the transcript, and two different colors reporting on transcription the 3’ end. In this scenario, the data would provide independent readouts of the same underlying signal, making it possible to perform two independent inferences on the same nucleus. This would allow for the decomposition of the inference into biological variability and inferential error using techniques analogous to those presented in S8.

Finally, we examined the inference method for spurious correlations to investigate the accuracy of the experimental single-cell correlations shown in Figure 5. The presence of spurious correlations would reflect inherent couplings in the inference method itself, since the simulation parameters were generated independently and stochastically.

Figure S5E-H show the single-cell correlations using the Spearman rank correlation coefficient between model parameters for the simulated dataset, as well as between the mean RNAP density and the cleavage time, as defined in the main text. Linear regression fits are also displayed for intuitive visualization. We discovered a slight positive correlation (ρ = 0.15) between the elongation rate and the cleavage time (Fig. S5G, *p* = 0.01). In contrast, there was no significant correlation between the mean initiation rate and the cleavage time, the mean initiation rate and the elongation rate, and the mean RNAP density and the cleavage time (Fig. S5E, F, and H). Although the relationship between the elongation rate and the cleavage time possessed the same, albeit weaker, correlation as found in the data (Fig. 5C), the main finding in the main text of the correlation between the mean RNAP density and the cleavage time was not reproduced by the simulations (Fig. S5H). The comparisons of Spearman rank correlation coefficients and *p*-values between the data and simulations are summarized in Table S2.

Thus, our results validated the single-cell correlations discovered in the main text, indicating that the experimental results were not the product of spurious correlations.

**Table S2.**
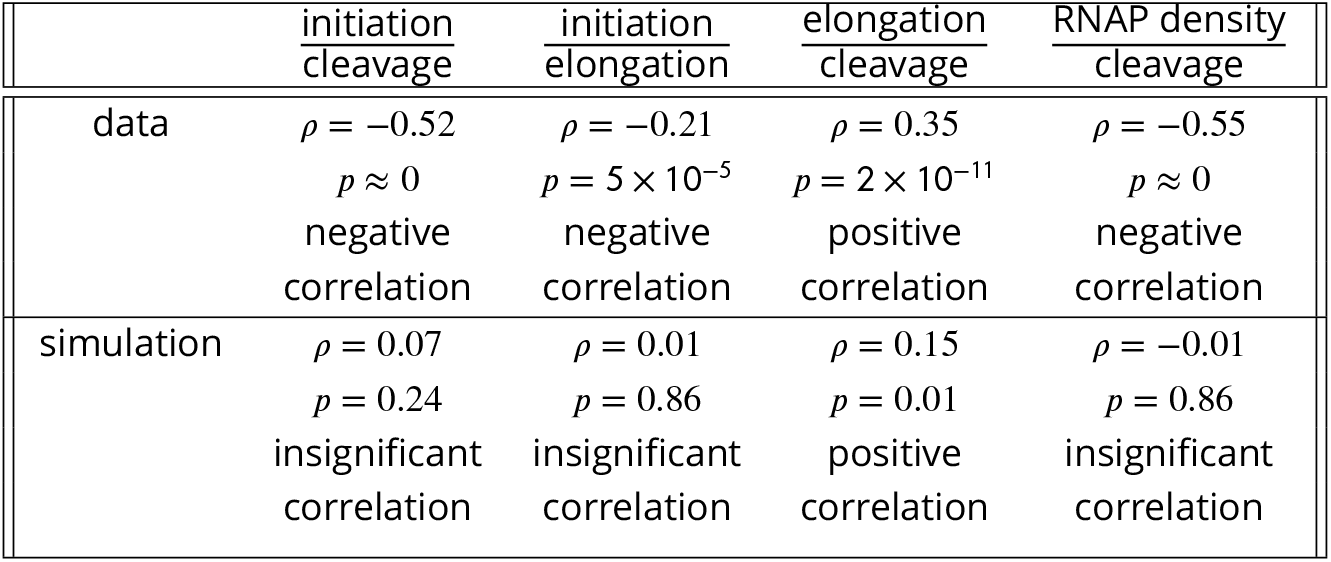
Comparison of Spearman rank correlation coefficients and *p*-values between experimental and simulated single-cell correlations.

|  | initiation<br>cleavage | initiation<br>elongation | elongation<br>cleavage | RNAP density<br>cleavage |
| --- | --- | --- | --- | --- |
| data | $\rho = -0.52$<br>$p \approx 0$<br>negative<br>correlation | $\rho = -0.21$<br>$p = 5 \times 10^{-5}$<br>negative<br>correlation | $\rho = 0.35$<br>$p = 2 \times 10^{-11}$<br>positive<br>correlation | $\rho = -0.55$<br>$p \approx 0$<br>negative<br>correlation |
| simulation | $\rho = 0.07$<br>$p = 0.24$<br>insignificant<br>correlation | $\rho = 0.01$<br>$p = 0.86$<br>insignificant<br>correlation | $\rho = 0.15$<br>$p = 0.01$<br>positive<br>correlation | $\rho = -0.01$<br>$p = 0.86$<br>insignificant<br>correlation |

## S5 Validation of the RNAP processivity assumption

The calibration between the MS2 and PP7 signals (Fig. 3) provided an opportunity to test the processivity assumption presented in the main text, namely that the majority of loaded RNAP molecules transcribe to the end of the gene without falling off. To estimate the processivity quantitatively, we assume that a series of *N* RNAP molecules transcribes past the MS2 stem loop sequence at the 5’ end of the reporter gene, and that only *pN* successfully transcribe past the PP7 stem loop sequence at the 3’ end. Here, we define *p* to be the processivity factor, and require 0 < *p* < 1. Thus, *p* = 1 indicates maximal processivity where every RNAP molecule that transcribes the MS2 sequence also transcribes the PP7 sequence, and *p* = 0 indicates minimal processivity, where no RNAP molecules make it to the PP7 sequence.

We assume that no RNAP molecules fall off the gene while they transcribe the interlaced MS2/PP7 loops used in the calibration experiment described in Figure 3B. Under this assumption, *N* RNAP molecules will fully transcribe both sets of stem loop sequences, allowing us to define the scaling factor as the ratio of total fluorescence values

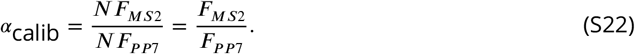

Note that, in this simple model, RNAP molecules can still fall off the gene after they transcribe the set of MS2/PP7 loops. Now, we consider the construct with MS2 and PP7 at opposite ends of the gene used in the main text. Allowing a fraction *p* of RNAP molecules to fall off the gene between the MS2 and PP7 loops, we arrive at a scaling factor

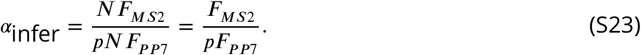

We can thus calculate the processivity *p* from taking the ratio of the true and biased scaling factors

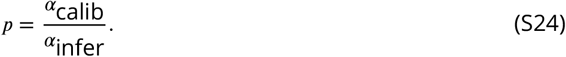

Taking the mean value of e*α*_calib_ from our control experiment using the interlaced MS2/PP7 loops to be the true value and the mean value of e*α*infer from the inference from the main text to be the biased value, we calculate a mean processivity of *p* = 0.96, with a negligible standard error of 4.81 × 10^−5^. Thus, on average, 96% of RNAP molecules that successfully transcribe the 5’ MS2 stem loop sequence also successfully transcribe the 3’ PP7 stem loop sequence, confirming previous results (***Garcia et al., 2013***) and lending support to the processivity assumption invoked in our model.

**Figure S6.**
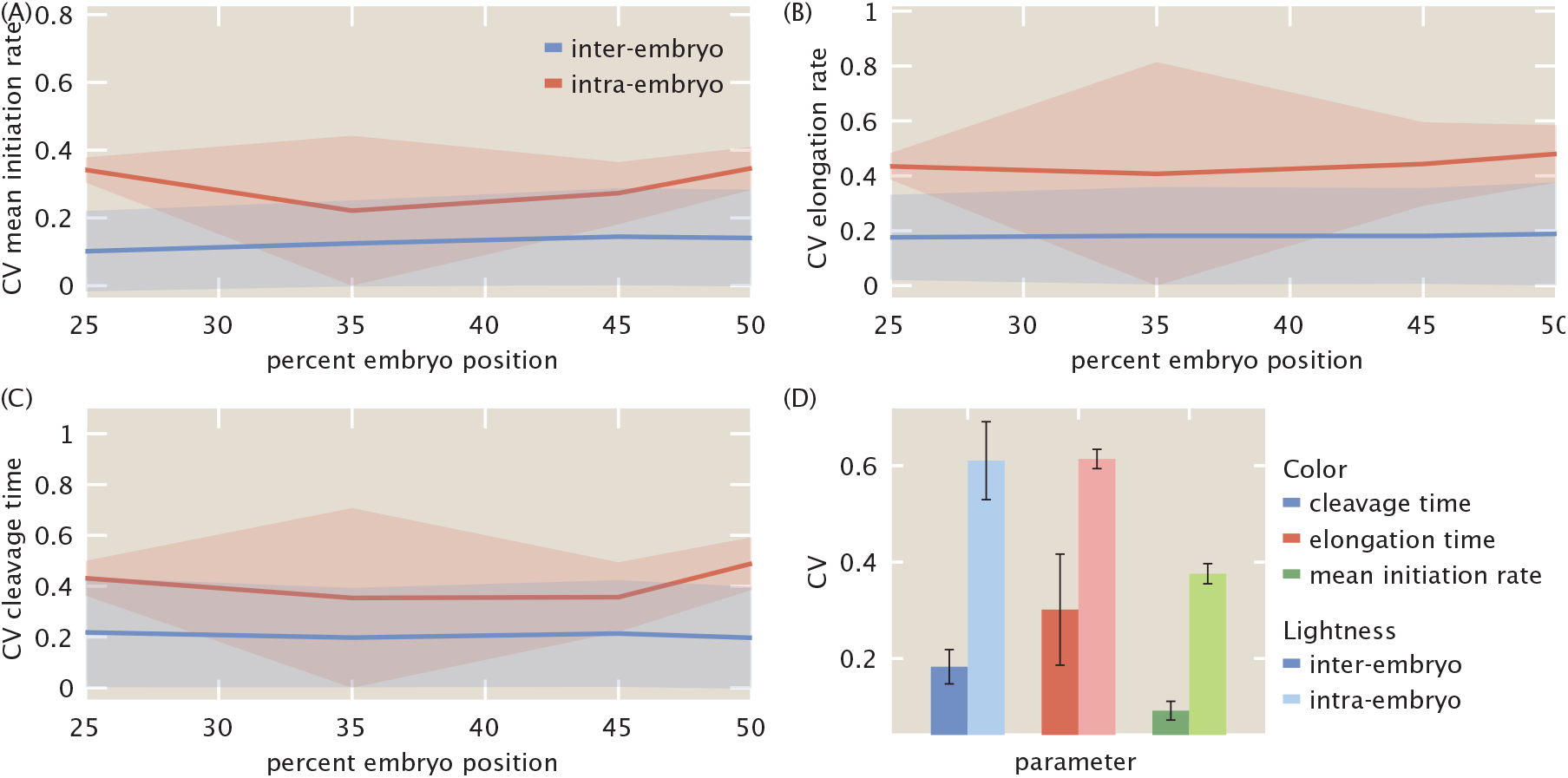
Comparison of intra- and inter-embryo variability for inferred (A) mean initiation rates, (B) elongation rates, and (C) cleavage times, as a function of embryo position. (D) Intra- and inter-embryo variability for transcriptional parameters averaged across all embryo positions. (A-C, lines and shaded regions indicate mean and standard error of the mean, respectively; D, error bars indicate bootstrapped standard error error across 100 bootstrap samples. Data were taken over 355 cells across 7 embryos, with approximately 10-90 cells per embryo in the region of the embryo examined here.)

## S6 Comparing intra- and inter-embryo variability

In the analysis in the main text, we treated all single cell inference results equally within one statistical set. In principle, this is justified only if the variability between single cells is at least as large as the variability between individual embryos. In this section we prove this assumption.

Here, we examine two quantities: the *intra-embryo variability*, defined as the variance in a parameter across all single cells in a single embryo, and the *inter-embryo variability*, defined as the variance across embryos in the single-embryo mean of a parameter. We examined these two quantities for the three primary inferred parameters—the mean initiation rate, elongation rate, and cleavage time.

Figure S6A-C shows the results of this comparison as a function of embryo position, where the red (blue) lines indicate the intra-(inter-) embryo variability and the red (blue) shaded regions indicate the standard error (bootstrapped standard error) in the intra-(inter-) embryo variability. For all of the parameters, the intra-embryo variability is at least as large as the inter-embryo variability, validating our treatment of all of the single-cell inference results as a single dataset, regardless of embryo.

This is seen more clearly when the data are averaged across embryo position. As shown in Fig. S6D, the inter-embryo variability of each parameter is substantially higher than the intra-embryo variability.

## S7 Full distributions of transcriptional parameters as a function of embryo position

Figure 4 presents inferred values of the transcriptional parameters in the form of population means and CVs as a function of embryo position. We chose this form of presentation to focus on spatial variation of these parameters via a succinct visualization.

**Figure S7.**
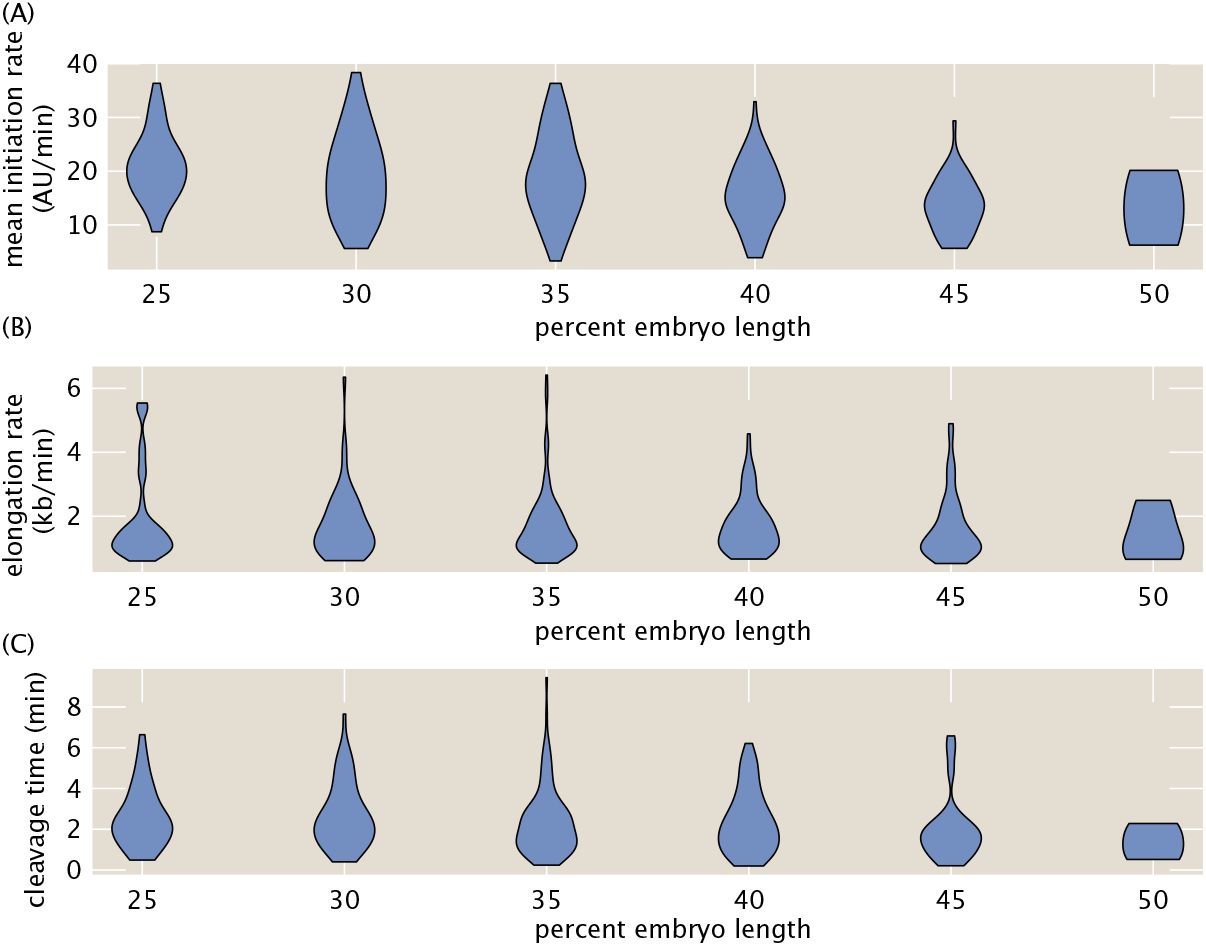
Single cell distributions of inferred parameters. (A-C) Full single-cell distributions of (A) mean initiation rate, (B) elongation rate, and (C) cleavage time as a function of embryo position.

Figure S7 shows the full distributions of the transcriptional parameters as a function of embryo position. For each parameter, the observed variability at a particular position in the embryo is quite broad, indicating substantial cell-to-cell variability. Nevertheless, there is no clear indication of multimodal behavior, indicating that the mean is still a reliable metric of population-averaged behavior.

## S8 Comparison of variability in mean initiation rate reported by our inference with static measurements

A widespread strategy to measure variability in transcription initiation relies on techniques such as single-molecule FISH (smFISH), which count the number of nascent transcripts at a transcribing locus in a fixed sample (***Femino et al., 1998***; ***Raj et al., 2006***; ***Zenklusen et al., 2008***; ***Little et al., 2013***; ***Zoller et al., 2018***). These single time point measurements are typically interpreted as reporting on the cell-to-cell variability in transcription initiation. Further, under the right conditions, the variability reported by this method has been shown to be dominated by biological sources of variability and to have a negligible contribution from experimental sources of noise (***Zoller et al., 2018***).

Inspired by these measurements in fixed embryos, we sought to determine how well our approach could report on biological variability. To do so, we contrasted the inference results of the transcriptional activity of our *hunchback* reporter with a snapshot-based analysis inspired by single-molecule FISH (***Zoller et al., 2018***). Specifically, we calculated the CVs in the raw MS2 and PP7 fluorescence in snapshots taken at 10 minutes after the start of nuclear cycle 14, from the same post-curation cells analyzed with the inference method. We reasoned that, since this calculation does not utilize the full time-resolved nature of the data, it provides a baseline measurement of total noise that encompasses both experimental and biological variability. As a point of comparison, we also calculated the CV in the instantaneous MS2 signal from another work using a similar P2P-MS2-lacZ construct (***Eck et al., 2020***).

Figure S8A shows the CV as a function of embryo position as reported by these different approaches. For the static measurements (red, green, and blue), the CV values lay around 20% to 80%. The CV of the inferred mean initiation rate (purple) exhibited similar values, although it was slightly lower in a systematic fashion. This difference was likely due to the fact that the inference relies on time-dependent measurements that can average out certain sources of error such as experimental noise, whereas such time averaging is not possible in the context of single time point measurements.

To succinctly quantify variability in the mean initiation rate, we then calculated the position-averaged squared CV for the same measurements in Figure S8A. The resulting squared CV values are shown in Figure S8B. Although the static measurements possessed essentially identical squared CVs (blue, red, green), the inference method exhibited a clear reduction in the squared CV (purple). To test whether the discrepancy in the variability between time-resolved and snapshot-based measurements arose from differences in the experimental error of each technique, we used the formalism introduced by ***Elowitz et al***. (***2002***) to separate the noise in the system into uncorrelated and correlated components. Here, uncorrelated noise represents random measurement error, while correlated noise contains both systematic measurement error as well as true biological variability. To perform this separation, we utilized the alternating MS2-PP7 reporter used in the calibration calculation (Fig. 3B). Because the MS2 and PP7 fluorescent signals in this reporter construct should, in principle, reflect the same underlying biological signal, deviations in each signal from each other should report on the relative magnitudes of both types of noise.

First, we defined the deviations δ_*MS*2_ and δ_*PP*7_ of each instantaneous MS2 and PP7 fluorescent signal from the mean MS2 and PP7 fluorescence signals, averaged across nuclei and time

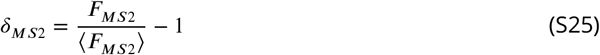

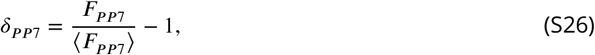

where *F*_*MS*2_ and *F*_*PP*7_ are the respective instantaneous MS2 and PP7 fluorescence values for a given nucleus and time point, and ⟨*F*_*MS*2_⟩ and ⟨*F*_*PP*7_⟩ are the respective mean MS2 and PP7 fluorescence values, averaged across nuclei and time points. Using these deviations, the uncorrelated and correlated noise terms are defined as

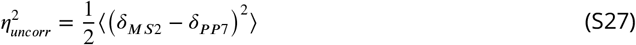

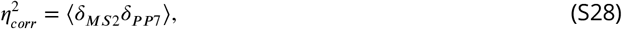

where the brackets indicate an ensemble average over time points and cells (***Elowitz et al., 2002***). From this, the total noise 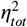, defined as the variance *σ*^2^ divided by the mean squared *μ*^2^, is simply the uncorrelated and correlated noise components added in quadrature

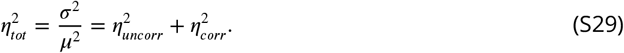

Note that the total noise 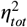 is simply the squared coefficient of variation. Thus, the squared coefficient of variation (CV^2^) of our data is equal to 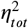 and can be separated into the uncorrelated and correlated components.

Figure S8B shows this CV^2^ (averaged across all embryo positions) for snapshots of the interlaced loop construct compared with the separated uncorrelated and correlated noise sources. Intriguingly, the uncorrelated and correlated noise (yellow) each contribute about half to the overall noise.

We posit that the relative magnitude of partitioning between correlated and uncorrelated noise also holds for the static measurements of spot fluorescence (Fig. S8B, blue, red and green). As a result, given this assumption, we can calculate the correlated and uncorrelated variability contributions to total squared CV from these static measurements. This is shown in light and dark red in the case of the static MS2 fluorescence measurement in Figure S8B. The figure reveals that the correlated noise component of the static measurements (dark red) is only slightly smaller than the overall noise measured by the inference (purple), suggesting that our inference method primarily reports on correlated variability.

**Figure S8.**
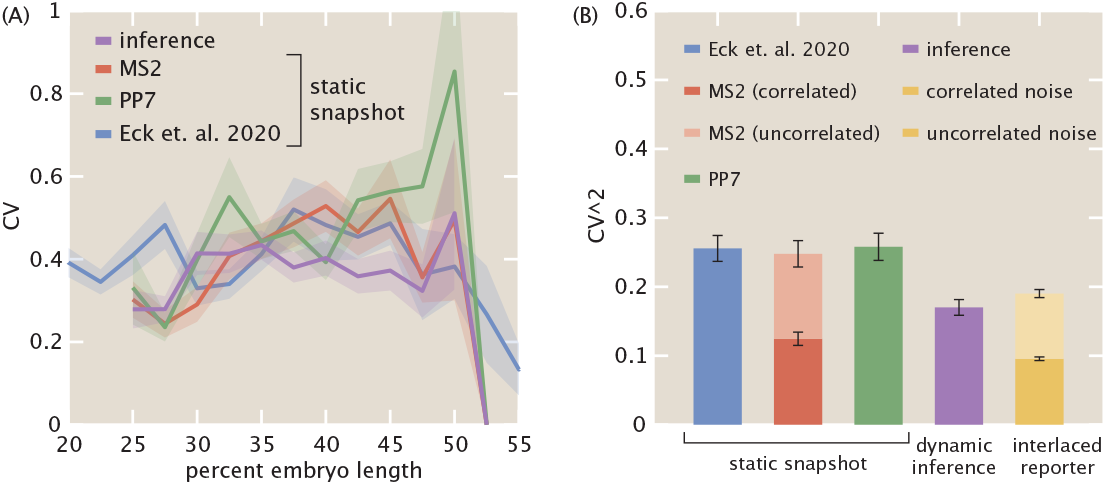
Comparison of coefficients of variation (CV) between inferred mean initiation rates and instantaneous counts of number of nascent RNA transcripts. (A) Position-dependent CV of inferred mean initiation rate (purple) compared with static measurements of MS2 and PP7 raw fluorescence (red, green) from the dual-color reporter (Fig. 1C), as well as with static measurements of MS2 data from ***Eck et al***. (***2020***) (blue). (B) Position-averaged squared CVs of the same measurements, where the entire dataset is treated as a single sample and embryo position information is disregarded. In addition, separation of uncorrelated and correlated sources of variability are shown, calculated using the reporter described in Fig. 3B. (A, Shaded regions indicate bootstrapped standard error of the mean; B, error bars indicate bootstrapped standard error of the mean for *n* = 100 bootstrap samples.)

As a result, the MCMC inference method can quantitatively capture the true biological variability in the mean initiation rate while separating out most of the uncorrelated contribution due to random experimental noise. Thus our results support the power of model-driven inference approaches in providing clean readouts of variability in transcriptional parameters.

## S9 Comparison of distribution of elongation rates with other works

As an additional validation of our inference results, we compared the distribution of single-cell inferred elongation rates with those reported in two similar works by ***Hocine et al***. (***2013***) and ***Fukaya et al***. (***2017***). Both of these works used a two-color live imaging reporter like the one utilized in this work, and measured the time delay between the onset of each stem loop signal to estimate a single-cell mean elongation rate. ***Fukaya et al***. (***2017***) studied a similar *hunchback* reporter to the one used here, while ***Hocine et al***. (***2013***) used a reporter construct in yeast.

Figure S9 shows the comparison of distributions of elongation rates. Because the reporter constructs and analysis techniques differed between works, a quantitative comparison is not possible. Nevertheless, all three sets of results report a significant cell-to-cell variability in mean elongation rate, ranging from 1 kb/min to 3 kb/min.

## S10 Theoretical investigation of single-cell distribution of elongation rates

To investigate the molecular mechanisms underling single-cell distributions of elongation rates obtained from the inference, we developed a single-molecule theoretical model. We were interested in how the observed variability in single-cell elongation rates could constrain models of the single-molecule variability in RNAP elongation rates. To disregard effects due to position-dependent modulations in the transcription initiation rate, we only studied cells anterior of 40% along the embryo length, where the initiation rate was roughly constant.

The model was adapted from the stochastic Monte Carlo simulation used in ***Klumpp and Hwa*** (***2008***), which accounts for the finite size of RNAP molecules (Fig. S10A). Here, single RNAP molecules are represented by one-dimensional objects of size *N*_*footprint*_ that traverse a gene consisting of a one-dimensional lattice with a total number of sites, corresponding to single base pairs, equal to *N*_*sites*_. The position of the active site of molecule *i* is given by *x*_*i*_, which takes integer values—each integer corresponds to a single base pair of the gene lattice. Because RNAP molecules have a finite size, given by *N*_*footprint*_, an RNAP molecule *i* thus occupies the lattice sites from *x*_*i*_ to *x*_*i*_ + *N*_*footprint*_. In this model, we do not incorporate sequence-dependent RNAP pausing along the gene.

**Figure S9.**
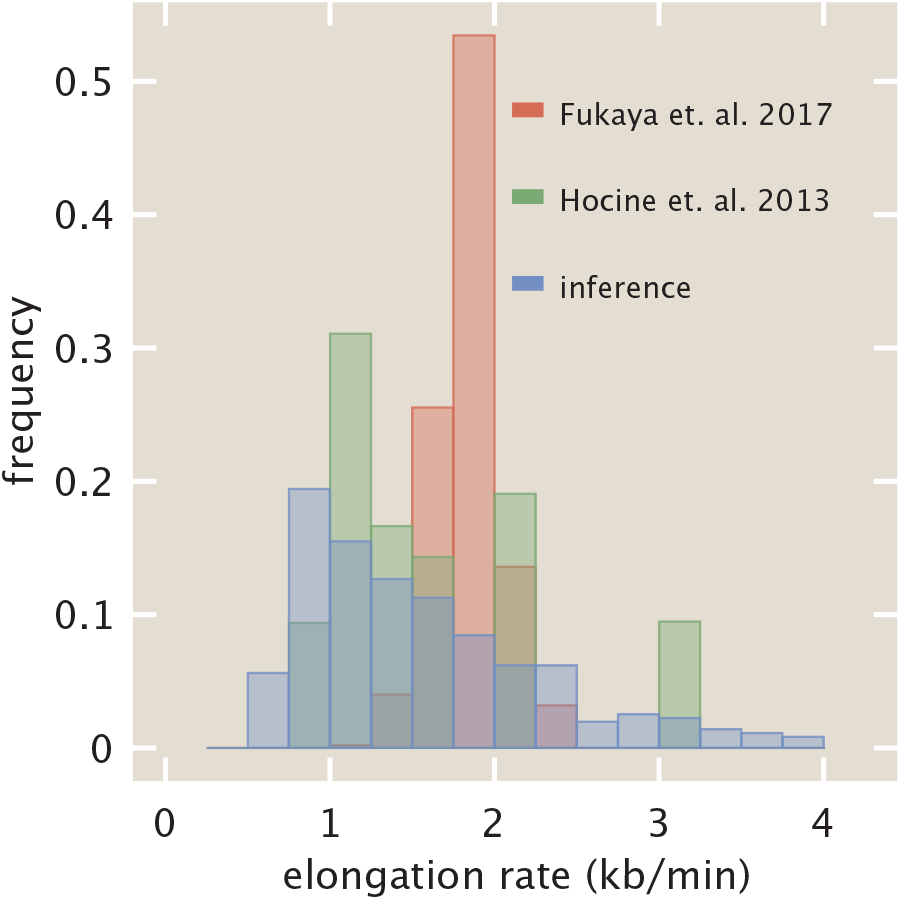
Comparison of distribution of elongation rates (green) with previous studies (***Hocine et al***. (***2013***), red and ***Fukaya et al***. (***2017***), blue). Distributions of previous studies were adapted from Figs. 2D and 2A of ***Hocine et al***. (***2013***) and ***Fukaya et al***. (***2017***), respectively.

New RNAP molecules are loaded at the start of the gene located at *x* = 0. Due to the exclusionary interactions between molecules, simultaneously simulating the motion of all molecules is unfeasible, and a simulation rule dictating the order of events is necessary.

At each simulation timestep *dt*, a randomized sequence of indices is created from the following sequence

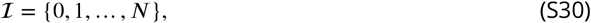

where {1, …, *N*} correspond to any RNAP molecules *i* = 1, …, *N* already existing on the gene, and 0 corresponds to the promoter loading site that generates new RNAP molecules.

Choosing indices *i* from the random sequence 𝔦 obtained above, the following actions are taken. If the index *i* indicates that an RNAP molecule was chosen (*i* > 0), then that RNAP molecule advances forward with stochastic rate *ε*. This probability is simulated by drawing a random number from a Poisson distribution with parameter *ε dt*, thus giving an expected distance traveled of *ε dt* per timestep (recall that, for a Poisson distribution with parameter *ε dt*, the resulting random variable corresponds to the number of occurrences in a time frame *dt*.). If this movement would cause the RNAP molecule to overlap with another RNAP molecule, then no action is taken. Otherwise, the RNAP molecule moves forward the number of steps given by the generated random variable.

If no RNAP molecule on the gene is chosen (*i* = 0), an RNAP molecule is loaded using a probability parameterized by the term *β dt*, only if no already existing RNAP molecules overlap with the footprint of the new RNAP molecule. If such an overlap occurs, then no action is taken. Otherwise, to calculate the probability of loading, a random number is drawn from a Poisson distribution with parameter *β dt*. If this number is one or higher, then the loading event is considered a success. The process is repeated until a total simulation time *T* has elapsed.

**Table S3.** Parameters used in single-molecule Monte Carlo simulation of elongation rates.

| Parameter | Description | Value |
| --- | --- | --- |
| $T$ | total simulation time | 600 sec |
| $dt$ | simulation timestep | 0.5 sec |
| $N_{sites}$ | size of lattice | 6626 bp |
| $N_{footprint}$ | RNAP footprint (Selby et al., 1997) | 40 bp |
| $\mu_\beta$ | mean loading rate | $0.17 \text{ sec}^{-1}$ |
| $\sigma_\beta$ | standard deviation of loading rate | $0.05 \text{ sec}^{-1}$ |
| $\mu_\tau$ | mean cleavage time | 2.5 min |
| $\sigma_\tau$ | standard deviation of cleavage time | 1.6 min |
| $\mu_\epsilon$ | mean elongation rate | free parameter |
| $\sigma_\epsilon$ | standard deviation of elongation rate | free parameter |

To simulate potential single-molecule variability, each RNAP molecule can possess a different stepping rate *ε*. For a given RNAP molecule *i*, its stochastic stepping rate *∈*_*i*_ is drawn from a truncated normal distribution *Tr* with mean *μ*_*∈*_ and standard deviation *σ*_*∈*_ and lower and upper limits 1 and infinity bp/sec, respectively

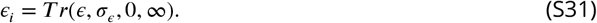

Once the position of the active site of an RNAP molecule exceeds that of the total number of sites *N*_*sites*_, i.e. the molecule reaches the end of the gene, it is removed from the simulation after the cleavage time *τ* elapses..

Finally, to account for single-cell variability in the transcription initiation rate, the loading rate *β* and cleavage time *τ* were allowed to vary across each simulated cell *j* by drawing these magnitudes from a Gaussian distribution with parameters reflecting the actual data. Since *hunchback* is known to load new nascent RNA transcripts at a rate of 1 molecule every 6 seconds in the anterior of the embryo (***Garcia et al., 2013***), we thus chose the mean of this distribution *μ*_*β*_ to be 1 molecule/6 *s* = 0.17 *s*^−1^. The standard deviation *σ*_*β*_ was chosen to be this mean multiplied by the CV in the initiation rate in the anterior inferred in the main text, resulting in a value of 0.05 *s*^−1^. Thus, for simulated cell *i*

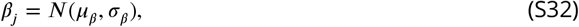

where any negative value was replaced with zero.

Similarly, the cleavage time *τ*_*j*_ for each simulated cell was drawn from a Gaussian distribution with mean *μ*_*τ*_ = 2.5 min and standard deviation *σ*_*τ*_ = 1.6 min. These values were obtained from the distribution of inferred cleavage times in the anterior of the embryo. The values of each simulation parameter are summarized in Table S3.

From these simulations, the positions of each RNAP molecule on the gene as a function of time were saved and then fed into the model of the reporter gene (Section S1), producing simulated single-cell MS2 and PP7 fluorescence traces (Fig. S10B). Simulated fluorescence noise was added using the same parameters as in the validation simulations discussed earlier (Section S5, Table S1, and Fig. S5). These fluorescence traces were then run through the inference pipeline (Section S4.1), resulting in inferred distributions of single-cell mean elongation rates from the single-molecule elongation simulation.

In order to compare these results with the empirically inferred distribution of elongation rates (Fig. 4D, red), we first considered a scenario where the single-molecule variability in stepping rates *σ*_*∈*_ was fixed at zero and the mean stepping rate *μ*_*∈*_ was varied from 0.6 to 2.1 kb/min. While the combination of exclusionary interactions between RNAP molecules, stochasticity in single-molecule stepping, and inferential noise did produce some cell-to-cell variability (Fig. S10C, top row), the resulting distributions nevertheless were unable to reproduce the large variance observed in the data. This can be seen by plotting the mean and variance of the simulated distributions (Fig. S10D, blue), where we see that the variance in the case of *σ*_*∈*_ = 0 is always below that of the data (Fig. S10D, purple).

Next, we allowed *σ*_*∈*_ to vary, simulating small to moderate variability with values of *σ*_*∈*_ = 0.3 kb/min and *σ*_*∈*_ = 0.6 kb/min. As expected, this single-molecule variability caused the inferred single-cell elongation rate distributions to widen (Fig. S10C, middle and bottom rows). In the presence of this variability, there existed parameter sets where the mean and variance of the simulated distributions quantitatively matched the empirical distribution within error (Fig. S10D, red and gold).

The distributions presented in the main text correspond to the following parameter values. For the case with no molecular variability in elongation rates (Fig. 4D, brown), we used *μ*_*∈*_ = 0.9 kb/min and *σ*_*∈*_ = 0 kb/min, chosen as the simulated parameter set with results closest to the inferred mean and variance of empirical elongation rates (Fig. S10D, lower black arrow). For the case with molecular variability in elongation rates (Fig. 4D, gold), we used *μ*_*∈*_ = 0.9 kb/min and *σ*_*∈*_ = 0.3 kb/min, chosen as a representative example of a simulation possessing a mean and variance in elongation rate that agreed with the inferred mean and variance of empirical elongation rates within error (Fig. S10D, upper black arrow), as well as qualitatively agreeing with the inferred distribution (Fig. 4D, gold).

## S11 Single-cell correlation analysis using full posterior distributions

The single-cell inter-parameter correlations presented in the main text (Fig. 5 were based off of mean values from the posterior distributions obtained from the inference procedure for ease of interpretation and visualization. In principle, these correlations could possess high amounts of uncertainty due to uncertainty in the single-cell parameter estimates. Here, we conduct a correlation analysis based on the full posterior distributions from the inference and validate the mean results presented in the main text.

To do so, we used a Monte Carlo simulation to construct a distribution of Spearman correlation coefficients and investigated if the mean Spearman correlation coefficients presented in Fig. 5 agreed with these simulated distributions.

First, we extracted the mean and variance of the inferred posterior distribution obtained from each single cell, for each transcriptional parameter (Fig. 2C and E). We then simulated *N* = 50, 000 new single-cell datasets comprising the mean initiation rate, elongation rate, and cleavage time, where these values were generated from Gaussian distributions parameterized by the means and variances from each parameter’s posterior distribution at the single-cell level.

Thus, each of the *N* = 50, 000 simulations resulted in a simulated dataset of *n* = 355 cells with randomly generated transcriptional parameter values obtained from the information inside the single-cell inferred posterior distributions from the experimental data. We then calculated an individual Spearman correlation coefficient and associated p-value for each simulation, generating an *N* = 50, 000 distribution for each correlation relationship.

Figure S11A and B show the ensuing distribution of p-values for the Spearman correlation coefficient between the mean initiation rate and elongation rate, as well as between the elongation rate and cleavage time, respectively. The p-values for the relationships between the mean initiation rate and cleavage time and between the mean RNAP density and cleavage time were essentially zero due to floating point error. Thus, the distributions of p-values for all four inter-parameter relationships were extremely small and support the statistical significance of their associated correlations.

Figure S11C shows the simulated distributions of Spearman correlation coefficients for all four relationships (histograms), along with the values obtained from the simpler mean analysis presented in the main text (dashed lines). We see that using the full posterior via this Monte Carlo simulation yields distributions that are in agreement with the results from the mean analysis, and that the distributions themselves are narrow, with widths of around 0.05. As a result, the correlations obtained from utilizing only mean inferred parameters quantitatively agree with the results obtained from utilizing the full Bayesian posterior obtained from the MCMC inference procedure.

**Figure S10.**
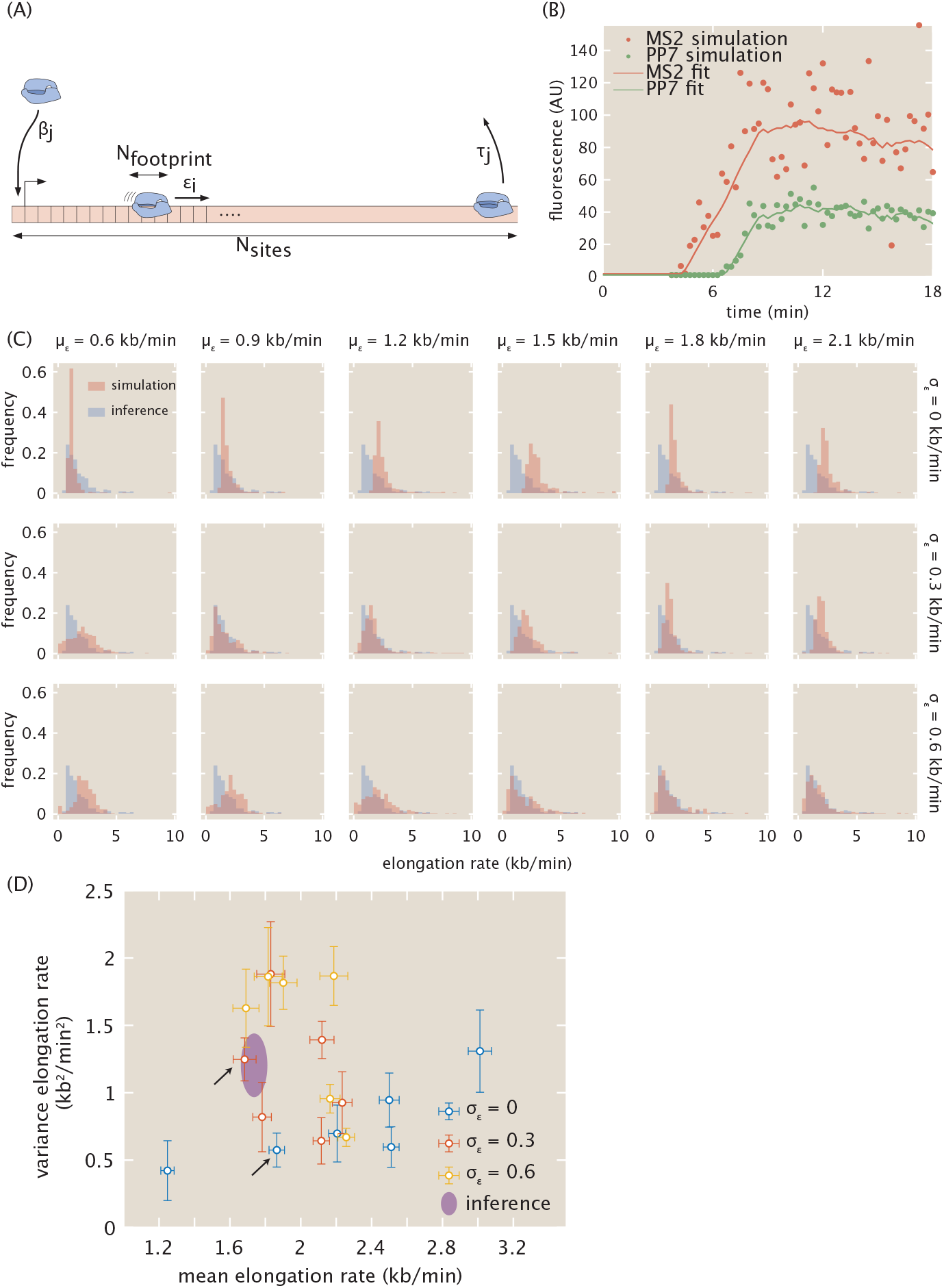
Single-molecule simulations of elongation dynamics require molecular variability to describe empirical distributions. (A) Cartoon overview of simulation. RNAP molecules with footprint *N*_*footprint*_ stochastically advance along a one-dimensional gene represented as a lattice with *N*_*sites*_ unique sites, with each site equivalent to a single base pair. Each RNAP molecule *i* possesses an intrinsic stepping rate *∈*_*i*_, and each cell *j* stochastically loads new RNAP molecules at the promoter with rate *β*_*j*_ and cleaves finished RNAP molecules after a cleavage time *τ*_*j*_. (B) Sample simulated MS2 and PP7 fluorescence traces for a single cell, using the single-molecule simulation with parameters *μ*_*∈*_ = 1.8 kb/min and *σ*_*∈*_ = 0 kb/min, along with inferred fits. (C) Simulated distributions of elongation rates (red) for varying values of *μ*_*∈*_ and *σ*_*∈*_, compared with inferred empirical distribution from data (blue). (D) Mean and variance of simulated and empirical distributions of elongation rates for varying values of *μ*_*∈*_ and *σ*_*∈*_. Without enough variability in the elongation rate of individual RNAP molecules (blue), the single-molecule model cannot produce the variance observed in the data (purple). However, in the presence of enough molecular variability, the empirical distribution’s mean and variance can be reproduced for certain parameter sets (red and gold). Black arrows correspond to parameter sets used for simulated distributions presented in the main text (Fig. 4D).

**Figure S11.**
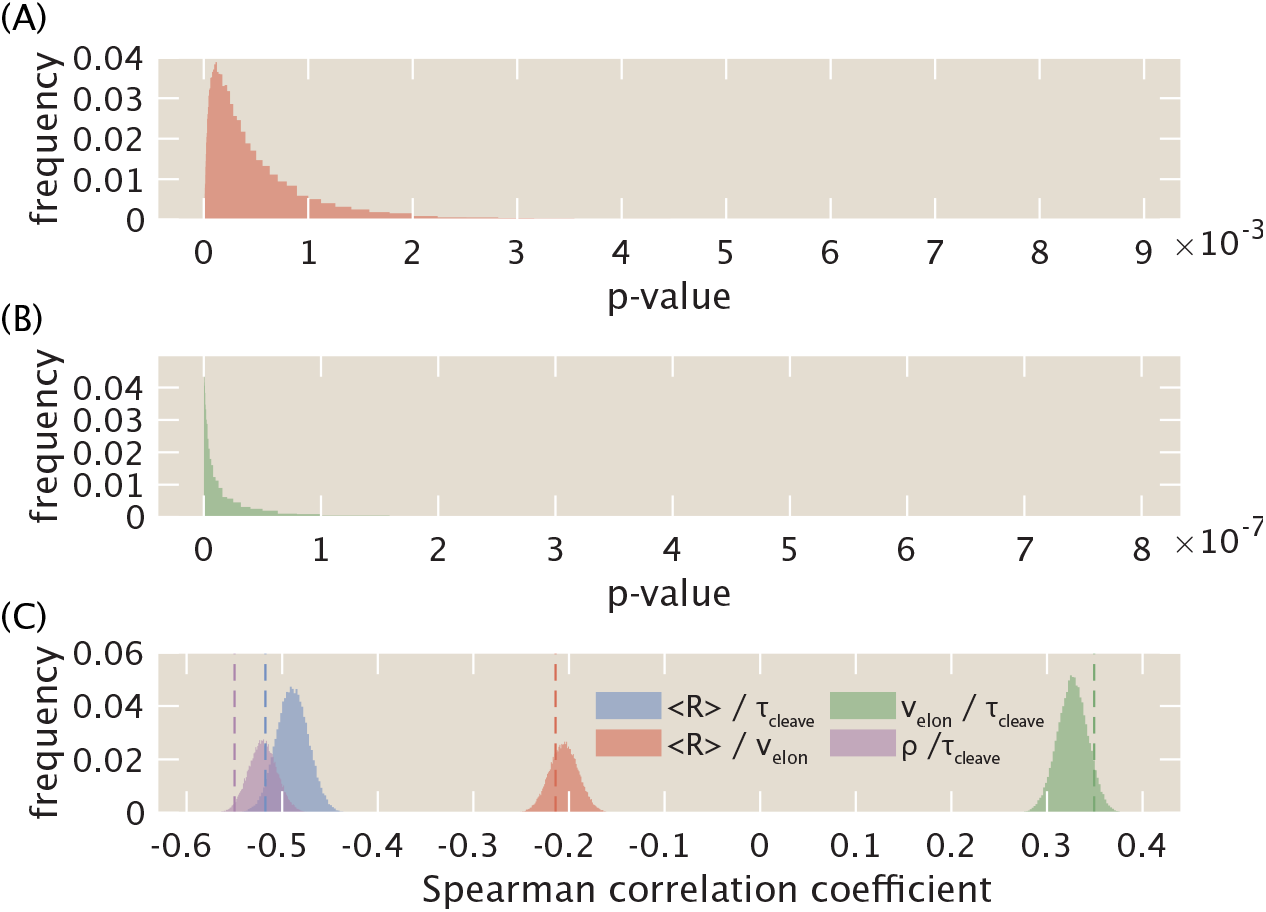
Monte Carlo simulation of error in single-cell analysis. (A, B) p-values of Spearman correlation coefficient for relationships between mean initiation rate and elongation rate (A) and between elongation rate and cleavage time (B). The p-values for the relationships between mean initiation rate and cleavage time as well as between mean RNAP density and cleavage time were essentially zero due to floating point error. (C). Distributions of Spearman correlation coefficients between mean initiation rate and cleavage time (blue), mean initiation rate and elongation rate (red), elongation rate and cleavage time (green), and mean RNAP density and cleavage time (purple). Results from mean-level analysis (Fig. 5) are shown in dashed lines.

Thus, our original analysis is robust, and we chose to retain its presentation in the main text for simplicity and ease of understanding.

## S12 Supplementary Videos

S1. **Video 1. Measurement of main reporter construct**. Movie of P2P-MS2-lacZ-PP7 reporter construct used in an embryo in nuclear cycle 14. Fluorescence intensities are maximum projections in the z-plane. Time is defined with respect to the previous anaphase.

S2. **Video 2. Measurement of interlaced reporter construct**. Movie of P2P-24x(MS2/PP7) reporter construct used in an embryo in nuclear cycle 14. Fluorescence intensities are maximum projections in the z-plane. Time is defined with respect to the previous anaphase.

